# Parietal cortex is recruited by frontal and cingulate areas to support action monitoring and updating during stopping

**DOI:** 10.1101/2025.02.28.640787

**Authors:** Jung Uk Kang, Layth Mattar, José Vergara, Victoria E. Gobo, Hernan G. Rey, Sarah R. Heilbronner, Andrew J. Watrous, Benjamin Y. Hayden, Sameer A. Sheth, Eleonora Bartoli

## Abstract

Recent evidence indicates that the intraparietal sulcus (IPS) may play a causal role in action stopping, potentially representing a novel neuromodulation target for inhibitory control dysfunctions. Here, we leverage intracranial recordings in human subjects to establish the timing and directionality of information flow between IPS and prefrontal and cingulate regions during action stopping. Prior to successful inhibition, information flows primarily from the inferior frontal gyrus (IFG), a critical inhibitory control node, to IPS. In contrast, during stopping errors the communication between IPS and IFG is lacking, and IPS is engaged by posterior cingulate cortex, an area outside of the classical inhibition network and typically associated with default mode. Anterior cingulate and orbitofrontal cortex also display performance-dependent connectivity with IPS. Our functional connectivity results provide direct electrophysiological evidence that IPS is recruited by frontal and anterior cingulate areas to support action plan monitoring/updating, and by posterior cingulate during control failures.

**In brief:** Functional connectivity between the intraparietal sulcus (IPS) and a set of frontal and cingulate regions indicates that IPS is recruited to aid inhibitory control. Control failures are associated with increased communication with posterior cingulate. IPS could be a novel and tractable neuromodulation target for control-related neuropsychiatric disorders.

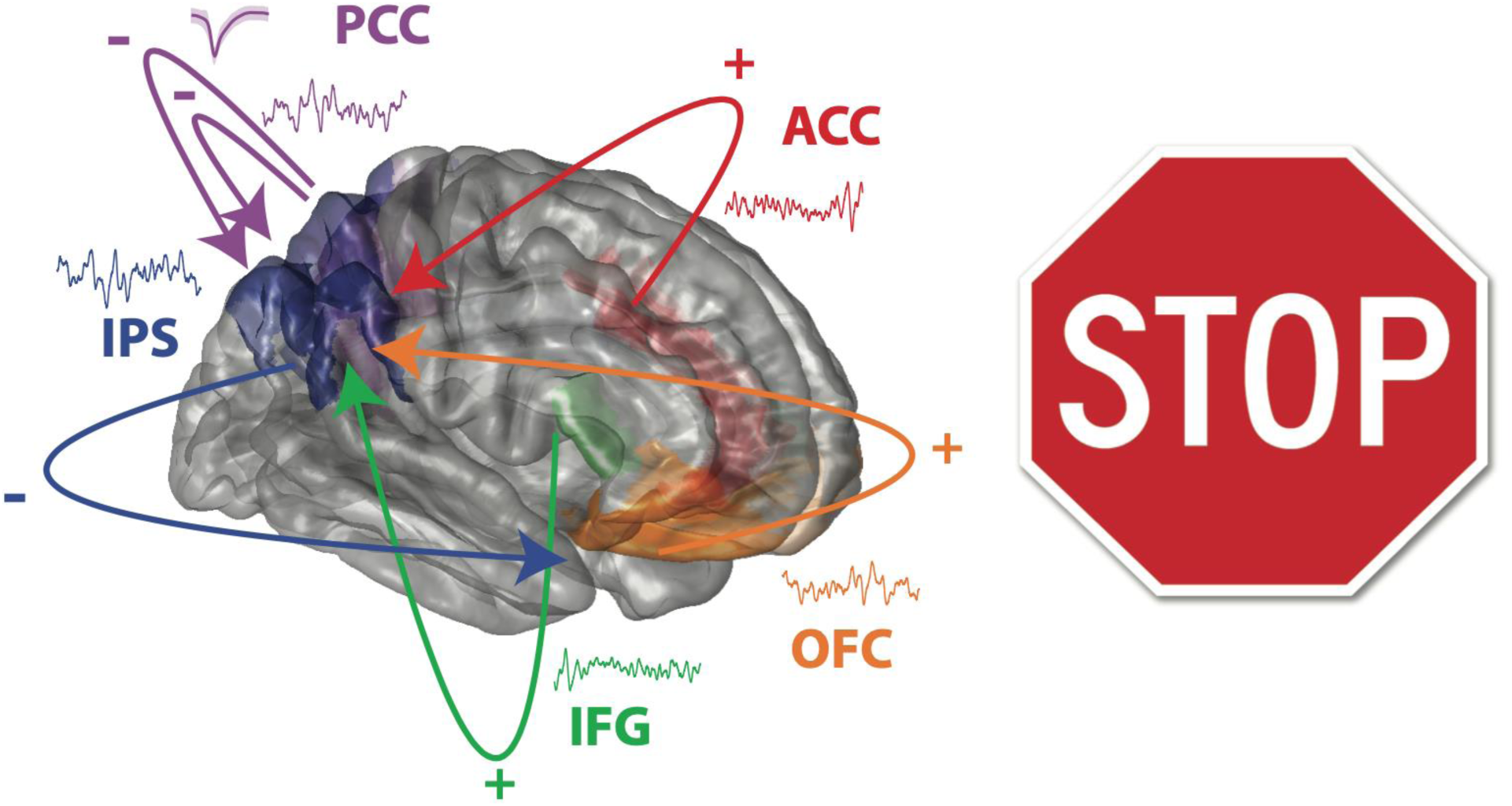

**Highlights:**

- Parietal cortex displays performance-dependent activity in action stopping
- Functional connectivity between IPS and IFG underlies successful stopping
- Early communication from ACC and OFC to IPS is also specific to successful stopping
- Communication from PCC to IPS is higher during lapses in control

## Introduction

Executive control manages the delicate balance between needs, contextual factors, priorities, and internal states to enable flexible and adaptive behaviors in our daily experiences.^1–3^ Stopping prepotent actions and automatic behaviors in response to changes in the environment is a crucial part of executive control.^4–8^ Suppression of unwanted or inappropriate actions can be broadly considered a key component for social interactions (e.g., language production or communication),^9–12^ and difficulties in this domain are associated with neurological conditions (e.g., tic or Tourette syndrome),^13–15^ and with psychiatric disorders related to impulsivity and lack of control (e.g., addictions, attention-deficit/hyperactivity disorder [ADHD], binge eating, obsessive-compulsive disorder [OCD], and post-traumatic stress disorder).^5,16–21^ Clinical trials aimed at treating control-related symptoms with non-invasive neuromodulation typically focus on prefrontal cortex.^22–24^ However, parietal regions may represent novel, unexplored targets to treat these conditions.

In laboratory settings, researchers employ the stop signal tasks (SSTs) to measure inhibitory control. In this paradigm, subjects perform fast-paced, repetitive actions and are required to withhold their response when a stop signal occurs, typically represented by a visual^25,26^ or auditory cue.^27^ A general consensus from previous human studies is that the right inferior frontal gyrus (IFG) starts the cascade of inhibitory control signals in the nervous system, then implemented through pre-supplementary motor area (pre-SMA) and the basal ganglia.^28–32^ Broadly, this type of inhibition is considered an subconstruct of cognitive control, typically associated with the frontoparietal and the cingulo-opercular networks.^33–36^ Indeed, several neuroimaging studies reported increased activity in the parietal cortex during SST,^37–43^ but its functional significance in inhibitory control, in contrast to frontal areas, historically received very little attention. Only a handful of recent studies directly investigated the role of parietal cortex during inhibitory control, with conflicting results.^18,37,38,44,45^ Transcranial magnetic stimulation (TMS) was employed to demonstrate that interfering with activity in the intraparietal sulcus (IPS) prolonged stop signal reaction times (SSRTs).^38^ This effect on behavior was specific to IPS, as TMS over the temporo-parietal junction (TPJ) did not produce any effect. Repetitive TMS results suggested the existence of parallel processing streams from ventral posterior IFG to pre-SMA and from dorsal posterior IFG to IPS.^37^ Overall, TMS-based evidence suggests that parietal cortex, specifically IPS, plays a critical role in inhibition, and these causal effects could be harmonized with the well-established parietal functions for spatial attention,^46–49^ reorienting,^50,51^ and adjusting movement planning.^52–54^ On the other hand, based on the latency of task-specific neural signals in the parietal cortex, occurring after SSRTs, others have concluded that IPS cannot play a causal role during inhibitory control.^18,44^ Inhibitory control signals are presumed to be transmitted sequentially from IFG to basal ganglia, motor cortex, and muscle, but not via IPS.^55^ However, the temporal order of the peak latency does not necessarily correspond to signal propagation in complex networks^56^ and the inertia to start and stop movements can smear temporal relations when we interpret neural signals with respect to behavioral measures,^57^ especially when these cannot be directly measured, like the SSRT.

To determine if IPS has the potential to be considered as a neuromodulation target for impulse control disorders, we need mechanistic evidence testing its recruitment with respect to the traditional set of regions important for inhibitory and cognitive control. To this end, we recorded intracranially from parietal, frontal and cingulate regions in human participants while they performed the SST. We leverage the high temporal precision afforded by intracranial recordings in human participants to test for the presence and direction of functional connectivity between IPS and other regions. If IPS plays a critical role in inhibition, we expect to detect activity and patterns of communication with IFG that depend on the success or failures in inhibitory control. We recruited 12 patients undergoing intracranial monitoring for epilepsy, and measured neural activity from the posterior parietal cortex (including IPS). Across this sample, we obtained simultaneous recordings from IFG and other brain regions that have been implicated in cognitive control:^33,58^ anterior cingulate cortex (ACC)^20,59–63^ and orbitofrontal cortex (OFC).^64–67^ We also recorded from posterior cingulate cortex (PCC), a region displaying a mixture of executive, default mode and memory-related functional properties across the literature.^68–72^ For both ACC and PCC, we further isolated single unit activity (SUA) to investigate signal dependencies with IPS at the neuronal level. Previous intracranial work primarily focused on single regions, and the few studies on inter-areal relations during inhibition did not focus on parietal cortex.^73,74^ We put forward the hypothesis that IPS and IFG will display patterns of communication in the beta frequency range (12-30 Hz) based on vast evidence of stopping-related effects in this band.^29,75–80^ If the role played by IPS is critical to the ability of successfully stopping a prepotent action, we expect two main results: a rich set of interconnections with the other regions and modulations in its communication patterns as a function of behavioral performance. In contrast, minimal interactions and/or lack of modulations depending on performance would indicate a limited role of IPS in inhibition, and thus very limited potential as a neuromodulation target for impulse control conditions.

## Results

We performed intracranial recordings of local field potentials (LFPs) from 12 patients while they performed a standard version of the SST (Figure 1A). For a subset of 6 patients, we also obtained extracellular measures of action potentials (SUA) in cingulate areas. Patients were recruited if they had at least one electrode recording from the posterior parietal cortex (i.e., parietal cortex excluding postcentral gyrus). The participants were required to press a keyboard button in response to the visual presentation of an arrow (“Go” trial), and to stop their response if the arrow changed color (“Stop” trial). The stop signal delay (SSD, delay between go and stop signal presentation) was adjusted adaptively based on performance to find the SSD value at which the participant failed at stopping 50% of the time (staircase method; see STAR Methods).

**Figure 1.**
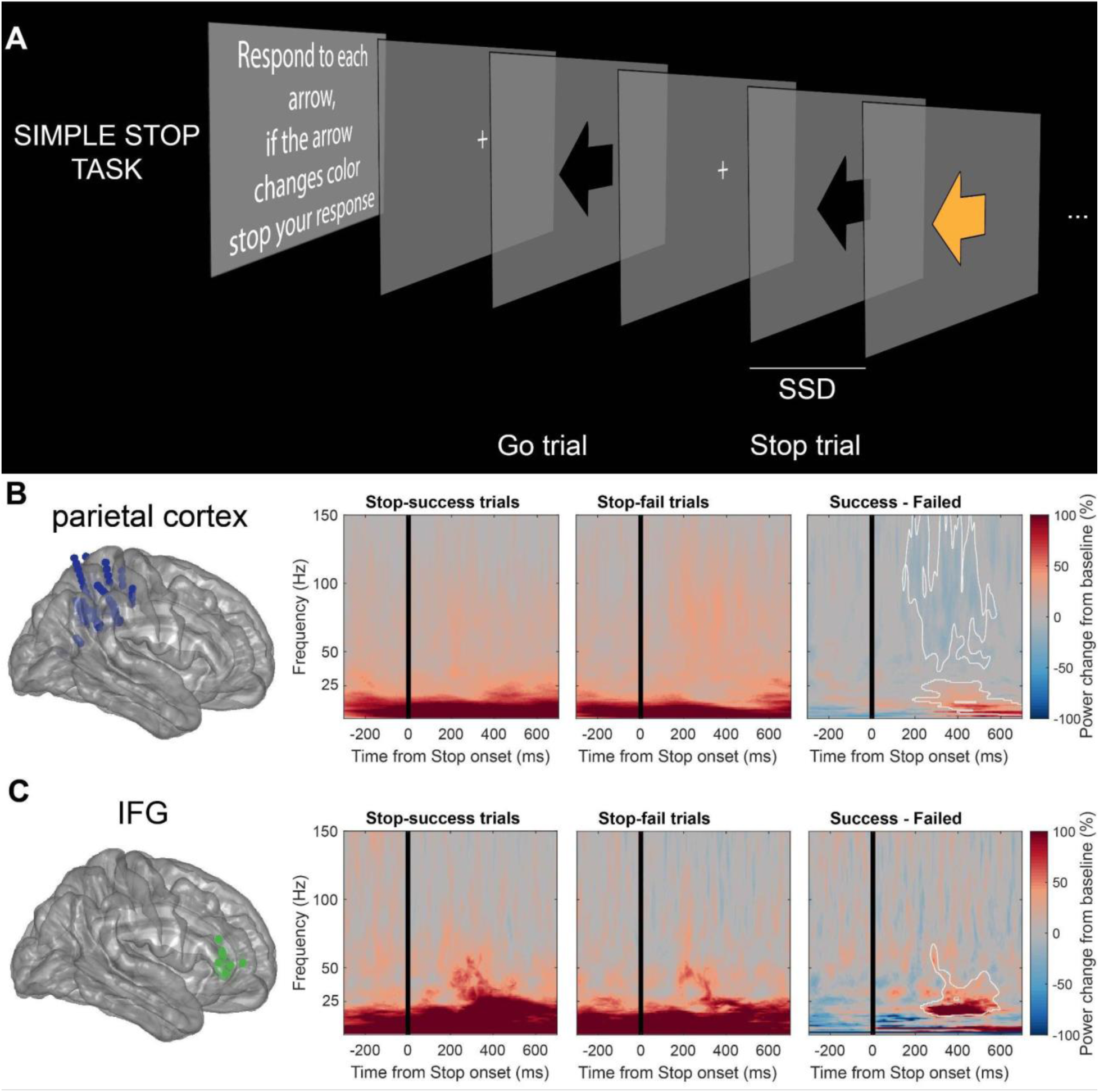
Experimental design. (A) Stop signal task. Each participant completed 288 trials: 216 Go trials (75 %) and 72 Stop trials (25 %). Stop signal delay (SSD) was initially set at 250 ms and adaptively varied so that each subject’s accuracy in stopping is 50%. Trial types were interleaved. See STAR Methods for details. (B-C) Time-frequency power (% change from baseline) averaged across recordings in (B) posterior parietal cortex (152 electrodes across 12 subjects) and (C) inferior frontal gyrus (27 electrodes across 6 subjects). From left to right, the time-frequency maps show the average values during Stop-success trials, during Stop-fail trials and their difference. Regions of significant differences (*p* < 0.05 corrected for multiple comparisons) are outlined in white on the difference map. Both parietal locations and IFG display higher power values during Stop-success trials starting around 200 ms after the stop signal (parietal: between 5-30 Hz; IFG: between 15-70 Hz). Parietal cortex additionally displays a decrease in broadband gamma values around the same time (spanning 35-150 Hz). Details for behavioral data and inclusion in time-frequency and connectivity analyses by each subject are shown in Table S1.

Performance measures and the estimation of the SSRT were computed following established procedures.^81,82^ Behavior across our sample displayed expected characteristics of the SST. The accuracy in Go trials was very high (median = 0.99, range 0.95 to 1.0). The probability of responding to a stop signal ranged between 0.33 and 0.51 (median = 0.47), demonstrating that the SSD adjustment achieved a balanced number of “Stop-success” trials (subjects successfully stopped their movements when a stop signal was presented) and “Stop-fail” trials (i.e., trials in which subjects failed to stop their movements and erroneously responded with a keyboard press). Reaction times were faster in Stop-fail trials than in Go trials (median = 676 ms versus 793 ms, Wilcoxon signed rank test, *p* < 0.001). The SSRT, estimated using the integration method, ranged between 120 and 330 ms across participants (median = 248 ms; see Table S1 in Supplemental Information).

First, we performed a time-frequency analysis of LFP power from all locations recording from the posterior parietal cortex, and compared power values associated with Stop-success and Stop-fail (permutation testing with cluster-based multiple comparison correction, *p* < 0.05) to evaluate if we could detect behaviorally-relevant differences in activity across parietal locations. Higher power values during Stop-success spanned low to mid frequency ranges (5 to 30 Hz), starting around 200 ms after the presentation of the stop signal and lasting about 500 ms (Figure 1B). Higher values for Stop-fail occurred across the broadband gamma range (35-150 Hz), similarly starting at about 200 ms after the stop-sign and lasting for 400 ms. This result demonstrated that the parietal cortex exhibits changes in neural activity with a timing that is compatible with the ongoing inhibition process, and with differences reflecting the success (or failure) of such inhibition. To offer a comparison with the well-established recruitment of IFG during stopping, we performed the same analysis on electrode locations recording from IFG, replicating previous evidence of an increase in beta power for Stop-success (significant cluster spanning 15-70 Hz) starting around 200 ms after the stop signal (Figure 1C). These results indicate the presence of a similar-timed response in both parietal cortex and IFG during stopping. To determine if the activity in these areas was a mere co-occurrence, or if it reflected actual neural communication, we investigated functional connectivity to and from a subdivision of posterior parietal cortex, IPS. For the connectivity analysis, we focused specifically on IPS based on previous evidence indicating a potential causal role of this specific parietal subdivision in inhibitory control.^38^

We measured functional connectivity to better quantify neural communication between IPS and IFG. We applied the same analysis between IPS and other regions (ACC, OFC, PCC). For these analyses, we employed a subsample of 7 patients (inclusion criteria: at least one electrode location in IPS). We used connectivity metrics between brain regions such as LFP-LFP Granger causality, LFP-LFP pairwise-phase consistency (PPC) and spike-LFP PPC.^83,84^ We specifically implemented LFP-LFP Granger causality and PPC because these two methods have been shown to co-vary reliably and provide us with a robust estimation of inter-areal communication.^85^ Spike-LFP measures reflect a complementary measure of neurophysiological interactions between brain regions based on the alignment of SUA in one region and the LFP in another region.^86–88^ Having consistent results across the three connectivity metrics will further increase confidence and generalizability of our results.^85,89^ With functional connectivity measures based on SUA and LFP, we inferred causal relationships between neural activity between different brain regions in the SST from signals at a much higher temporal resolution than previous fMRI studies. Overall, our results show that IPS is primarily a recipient rather than a sender of information from IFG and other regions during action stopping, with rich interconnections and modulations related to the behavioral outcome, detailed in the following sections. This is consistent with IPS being recruited to aid movement inhibition and playing a critical role in the ability to successfully inhibit an ongoing action.

### Information primarily flows from other brain regions to IPS in successful movement inhibition

We quantified direction and strength of information flow between IPS and other brain regions using LFP-LFP spectral Granger causality and PPC in three different but partially overlapping epochs in Stop trials aligned to the stop signal onset: [-800 ms 0 ms] “Before-stop”; [-400 ms +400 ms] “During-stop”; and [0 ms +800 ms] “After-stop”. With this approach, we aimed to assay how information flow between brain regions subserves stopping behavior in preparation (Before-stop), in inhibition (During-stop), and after stopping (After-stop). We estimate Granger causality across a wide spectral range (10-128 Hz), and we are interested in assessing if the interactions between areas are more prevalent in the beta frequency range, given the prominent role of this frequency band in movement inhibition.^6,75,90^

First, we focus on LFP-LFP spectral Granger causality between IPS and IFG. We use “Granger causality from A to B” as a simplified terminology to describe how LFP from area B can be effectively predicted based on LFP from area A. We measured Granger causality in the three different 800 ms intervals aligned to the onset of the stop signals (Figure 2). In the Before-stop epoch, Granger causality from IFG to IPS was significantly higher than vice versa at 23-26 Hz (Wilcoxon signed rank test, *p* < 0.01 at each frequency). Peak IFG-to-IPS Granger causality is 0.021 at 16 Hz, and peak IPS-to-IFG Granger causality is 0.015 at 20 Hz. From the chance level of 0.007, this is a ratio of 1.8:1 (Figure 2A). This demonstrates the presence of an early influence of IFG over IPS. There was no significant difference in Granger causality from IPS to IFG and from IFG to IPS in the During-stop epoch (Figure 2B). The lack of difference does not correspond to a lack of communication: as evident in the figure, the Granger causality values are higher than chance (represented by the horizontal gray shaded area) and thus are consistent with symmetric exchange of information between the two areas. For simplicity, we refrain from focusing on symmetric information flow in the Granger causality results, as our main objective with this metric is to identify directionality effects. In the After-stop epoch, Granger causality from IPS to IFG was significantly higher than vice versa at 13-16 and 31-36 Hz (Wilcoxon signed rank test, *p* < 0.01 at each frequency). Peak IPS-to-IFG Granger causality is 0.017 at 14 Hz, and peak IFG-to-IPS Granger causality is 0.012 at 23 Hz. From the chance level of 0.007, this is a ratio of 2:1 (Figure 2C). Crucially, the asymmetry in Granger causality is not due to signal-to-noise ratios (Figure S1).^83,91^ Granger causality results in Stop-success trials were similar when we limit our analyses in right IPS and right IFG only (Figure S2). There was no significant asymmetry in information flow between IPS and IFG across the three different epochs in Stop-fail trials (Figures 2D-F). However, when considering the right hemisphere only, there was asymmetry in Granger causality between right IFG and right IPS in Stop-fail trials (Figures S2D-F): IFG-IPS asymmetry in Before-stop epoch (similar to Stop-success) followed by IPS-IFG asymmetry in the During-stop (different from Stop-success). Thus, functional connectivity between IPS and IFG shows modulations based on task outcomes. Patterns associated with Stop-success were highly preserved within hemispheres, while Stop-fail exhibited more variable patterns, but in any case distinct from those underlying correct inhibition.

**Figure 2.**
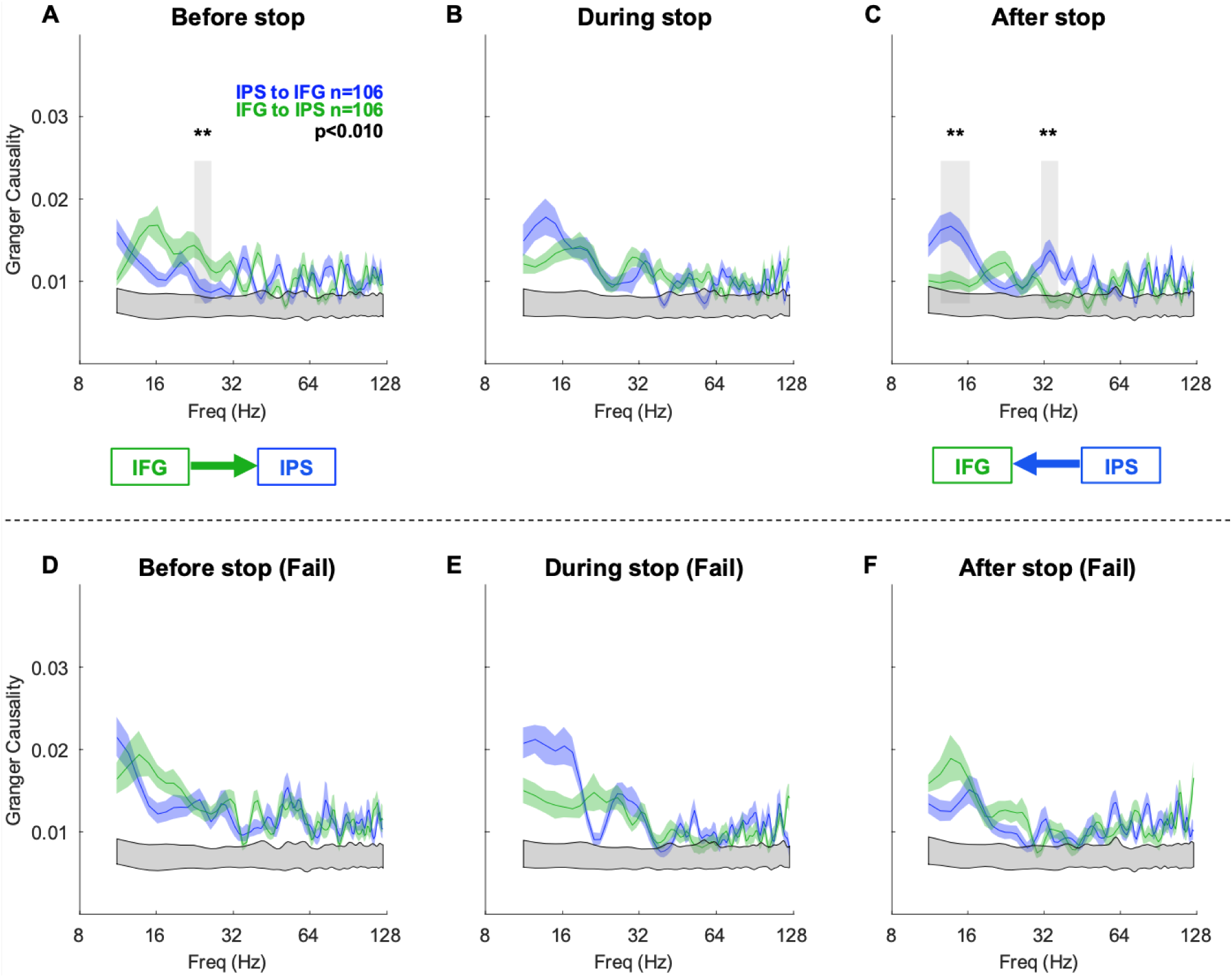
Granger causality between IPS and IFG in the SST across different time periods in movement inhibition. We measured Granger causality from IPS to IFG (blue) and from IFG to IPS (green) in the epochs aligned to the Stop Cue presentation as the following: (A) [-800 ms 0] “Before-stop”; (B) [-400 ms +400 ms] “During-stop”; and (C) [0 +800 ms] “After-stop”. In all panels, the black asterisks and gray vertical shaded regions denote *p* < 0.01 (Wilcoxon signed-rank test); Gray shaded regions in the bottom are the 99% bounds of a permutation test (see STAR Methods); Colored traces denote mean Granger causality values; Colored shaded regions denote ± standard error of the mean (SEM); n indicates the number of LFP-LFP pairs. (A) In the Before-stop epoch, Granger causality is significantly higher from IFG to IPS than vice versa at 23-26 Hz. The schematic summarizes that there is a significantly asymmetric communication from IFG to IPS. (B) In the During-stop epoch, there is no significant difference between Granger causality from IFG to IPS and vice versa. (C) In the After-stop epoch, Granger causality is significantly higher from IPS to IFG than vice versa at 13-16 and 31- 36 Hz. The schematic summarizes that there is a significant asymmetric communication from IPS to IFG. (D-F) There is no significant difference between IPS-to-IFG and IFG-to-IPS Granger causality in Stop-fail trials. Formats are the same as (A-C).

We also assayed information flow between IPS and the other brain regions (ACC, OFC, PCC) using Granger causality (Figures 3, S3, and S4). In Stop-success trials, OFC sends more information to IPS than vice versa in the Before-stop epoch (13-18 Hz; Figures 3A and S3A), mirroring the result obtained for IFG. In the During-stop, the communication reversed (IPS to OFC) and in the After-stop epoch there was no significant asymmetry in communication between OFC and IPS (Figures 3B-C and S3B-C). The directionality of communication between IPS and ACC and between IPS and PCC does not change significantly across the three different epochs. ACC sends more information to IPS than vice versa in low gamma (35-64 Hz) and high gamma (64-128 Hz) frequency range (Figures 3A-C and S3D-F) while PCC and IPS did not show any directionality (i.e, symmetric exchange of information) in all three different epochs (Figures 3A-C and S3G-I).

**Figure 3.**
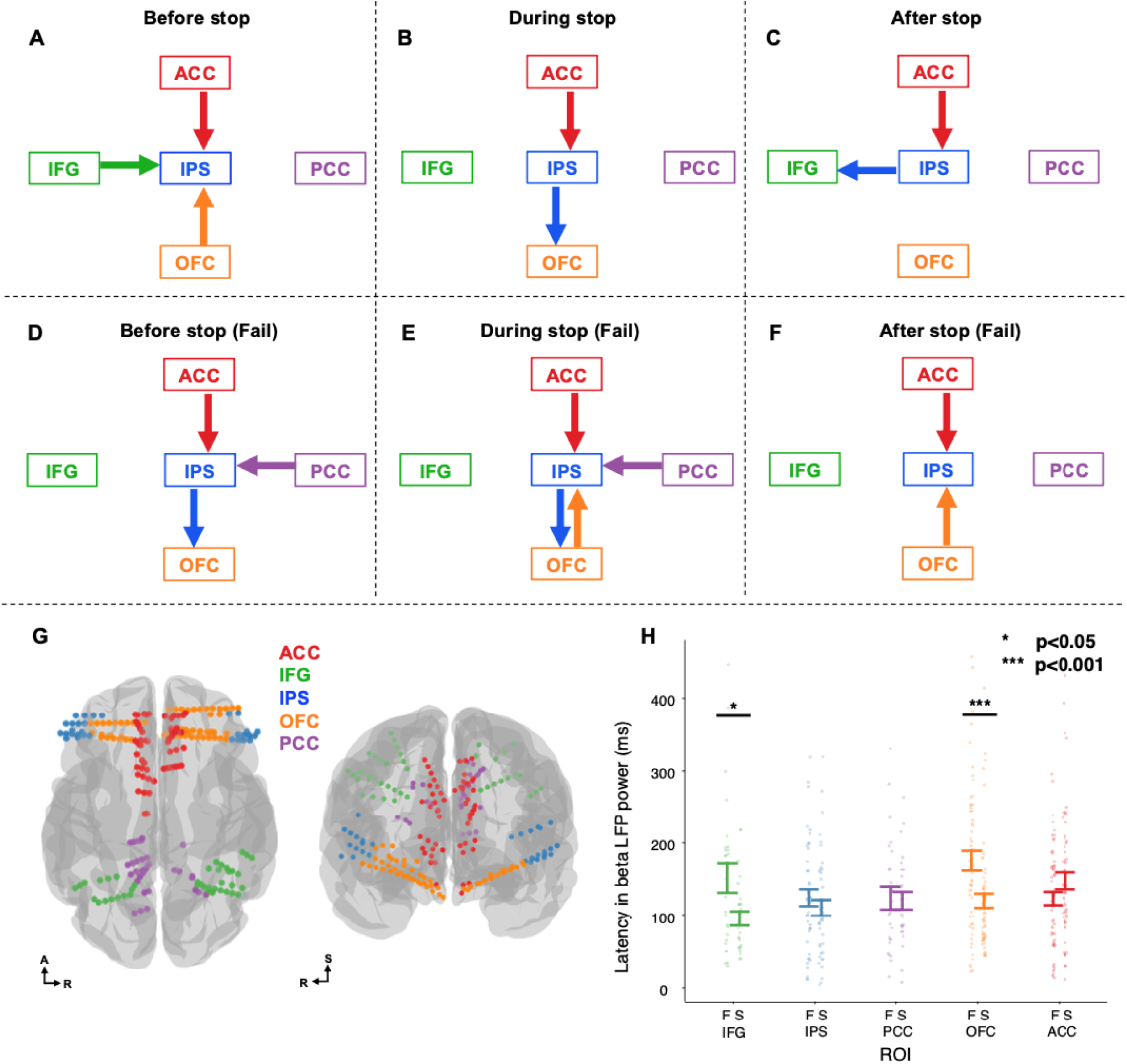
Beta band LFP latency and Granger causality results from IPS and other brain regions of interest. (A-C) A schematic of information flow based on Granger causality in Stop-success trials. An arrow indicates significant asymmetric information flow between the two brain regions with the larger information flow in the arrow direction. No arrow is consistent with symmetric exchange of information flow. Frequency information is not displayed for simplicity. Details are shown in Figure S3. (A) In the Before-stop epoch, information primarily flows from IFG, OFC, and ACC to IPS. (B) In the During-stop epoch, information primarily flows from ACC to IPS and from IPS to OFC. (C) In the After-stop epoch, information primarily flows from IPS to IFG and from ACC to IPS. (D-F) A schematic of information flow based on Granger causality in Stop-fail trials. Formats are the same as (B-D). Details are shown in Figure S4. (D) In the Before-stop epoch, information primarily flows from ACC and PCC to IPS and from IPS to OFC. (E) In the During-stop epoch, information primarily flows from ACC and PCC to IPS with asymmetric exchange of information between IPS and OFC in different frequency ranges. (F) In the After-stop epoch, information primarily flows from ACC and OFC to IPS. (G) Electrode locations employed in the Granger causality analysis between IPS (n=41) and all other regions: IFG (n=27), ACC (n=57), OFC (n=58), and PCC (n=25) displayed in MNI coordinates on a brain template (fsaverage). Electrode locations for each individual subject are shown in Figure S5. (H) First peak timing in LFP beta (15-25 Hz) in ACC, IFG, IPS, OFC, PCC. Mean latency of the beta power peak (from the stop signal presentation) and standard error for each region divided by Stop-success trials, denoted by “S” and Stop-fail trials, denoted by “F”. Each dot displays the data from each electrode location. There was a significant performance-specific modulation in IPS (*p* < 0.05) and OFC (*p* < 0.001) based on the mixed-effects model.

In Stop-fail trials, however, we observed different patterns of information flow between IPS and the other brain regions. In the Before-stop epoch, there was higher communication from IPS to OFC, and in the During-stop epoch there were different peaks in the two directions (peak IPS-to-OFC Granger causality 0.021 at 25 Hz; peak OFC-to-IPS Granger causality 0.021 at 13 Hz; Figures 3D-E and S4A-B). In the After-stop epoch, Granger causality from OFC to IPS was significantly higher than vice versa, which was not observed in Stop-success trials (Figures 3F and S4C).^92^ With respect to ACC, in addition to the modulation in low and high gamma frequency that was already observed in Stop-success trials (Figures 3A-C and S3D-F), Granger causality from ACC to IPS in Stop-fail trials is significantly higher than vice versa at 21-26 Hz in the After-stop epoch (Figure S4F). Interestingly, Granger causality from PCC to IPS was higher at 16-20 Hz in the Before-stop epoch (Figures 3D and S4G) and at 50-59 Hz in the During-stop epoch (Figures 3E and S4H). Electrode locations used for the analysis are displayed on a template (Figure 3G). Overall, failures in inhibition were associated with stronger interactions between IPS and OFC and more communication from ACC to IPS in the After-stop epoch. Communication from PCC to IPS was specific to failures in inhibition, not being present during Stop-success.

### Latency in LFP power in beta frequency shows modulations based on stop success and failure in IFG and OFC

To further test the temporal relationship of activity between brain regions, we compared the latency of LFP signals in both beta and gamma frequency ranges for Stop-success and Stop-fail trials in each region. We computed the latency of the first peaks in beta and gamma frequency (same patients and electrodes used for LFP-LFP connectivity analyses, Table S2 and Figure S8). Considering beta-band, the mixed-effects model showed a significant difference between the trial types, with the beta peak occurring earlier for Stop-success versus Stop-fail (*p* = 0.013, mean and SEM: Success = 123±29 ms; Fail: 142±34 ms). No overall effect of region was evident (*p* = 0.32), while the interaction between region and trial type was significant (*p* = 0.0013, Figure 3H). Post-hoc testing revealed that beta activity in IFG had earlier first peaks in Stop-success trials than Stop-fail trials (Success: 96±19 ms; Fail: 152±40 ms; corrected *p* = 0.011). The same pattern was true for OFC (Success: 121±29 ms; Fail: 175±39 ms; corrected p=0.0003). IPS showed a similar modulation, but it did not reach significance in the post-hoc analysis (Success: 110±28 ms; Fail: 124±29 ms; p=0.43). PCC showed no modulation (Success: 120±23 ms; Fail: 124±30 ms; p=0.87), while ACC displayed a trend in the opposite direction, with Stop-fail trials displaying an earlier peak (Success: 148±33 ms; Fail: 123±26 ms; p=0.09). The analysis was repeated for the broadband gamma range, finding no overall effects for trial type, region, or their interaction (all p-values >0.05). The overall delay in the IFG beta peak associated with failed inhibition is compatible with a delay in the inhibition process, starting the inhibition process later than the go process, in line with race model predictions. Overall, these results also demonstrate that IFG shows the earliest beta-band peak, followed by IPS, PCC, OFC and ACC.

### Information flow between parietal cortex and neurons in PCC is higher during fail trials

We collected SUA in ACC and PCC. We did not have any recordings of SUA from the other regions (IFG, IPS, OFC) as they are not typical targets for our microwire probes. We quantified spike-LFP coherence by using SUA in ACC and PCC and evaluating the pairwise phase-consistency of spike-timing with respect to beta range LFP in parietal cortex (see STAR Methods).^93^ We use “spike-LFP PPC” or “spike-LFP coherence” as a simplified terminology to describe how SUA in ACC or PCC is coherent with LFP phase in the beta range measured from parietal cortex. Similar to LFP-LFP Granger causality results, spike-LFP coherence between ACC and parietal cortex showed no modulation based on Stop-success and Stop-fail trials (Figures 4A and 4C, red; paired t-test, *p* > 0.1). Spike-LFP coherence between PCC and parietal cortex was higher in Stop-fail trials than in Stop-success trials (Figures 4B and 4C, purple; paired t-test, *p* < 0.05), and this result is consistent with the pattern observed in LFP-LFP Granger causality (Figures 3 and S4G-I). Finding a performance-specific information flow from PCC to parietal cortex at the neuronal level further supports the notion that the communication between PCC and parietal cortex is specific to failures in control.

**Figure 4.**
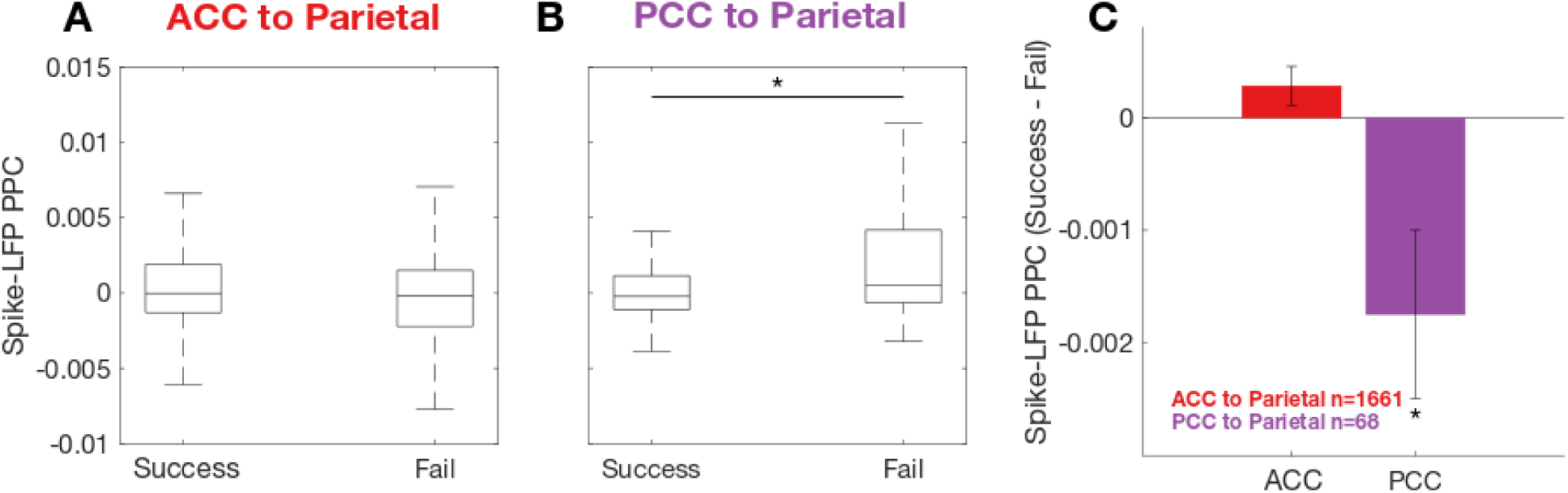
Spike-LFP PPC in 15-25 Hz from ACC to parietal cortex and from PCC to parietal cortex. We measured spike-LFP PPC in Stop-success and Stop-fail trials. For each sorted unit in a given participant, we computed spike-LFP PPC across all LFP signals in parietal cortex in that participant. Spike-LFP PPC from ACC to IPS is shown in red. (A) A box plot of all PPC values based on individual spike-LFP pairs from ACC to parietal cortex (n=1661) in Stop-success and Stop-fail trials. Outliers are omitted. (B) A box plot of all PPC values based on individual spike-LFP pairs from PCC to parietal cortex (n=68) in Stop-success and Stop-fail trials. Spike-LFP PPC values are significantly higher in Stop-fail trials than in Stop-success trials (paired t-test, *p* < 0.05). (C) Difference in PPC values between the two trial types. Positive difference indicates higher communication during Stop-success trials, and negative difference indicates higher communication during Stop-fail trials. There was no significant modulation based on success or fail trials in spike-LFP PPC from ACC to parietal cortex (paired t-test, *p* > 0.1). Spike-LFP PPC from PCC to parietal cortex is shown in purple. Spike-LFP PPC from PCC to parietal cortex was significantly higher in Stop-fail trials than in Stop-success trials (paired t-test, *p* < 0.05). Error bars denote ±SEM.

### Bidirectional information flow between IPS and other brain regions is higher during success than fail trials

Next, we used LFP-LFP PPC to measure phase alignment, an undirected metric of connectivity, between IPS and the other brain regions. With Granger causality, we could infer both direction and strength of communication. The goal of Granger causality analysis was to infer which brain region was a sender and therefore a driver of communication by comparing the incoming and outgoing Granger causality values. In contrast, with PPC we specifically measure bidirectional communication, quantifying whether the signals from two regions display a consistent phase alignment at each frequency, regardless of their amplitude modulations. To test whether phase-based bidirectional connectivity between two brain regions subserves motor inhibition, we measured the difference in LFP-LFP PPC in Stop-success and Stop-fail trials in the same three epochs that we used for Granger causality: Before-stop, During-stop, and After-stop. Positive values in the difference in PPC are consistent with an increased phase alignment during successful motor inhibition, negative values with a consistent phase alignment during stopping failures. Our results in LFP-LFP PPC analysis showed that bidirectional information flow between IPS and IFG (Figures 5B), IPS and OFC (Figures 5D and S7C), and IPS and ACC (Figures 5D and S7D) is important for stopping behavior, detailed below.

**Figure 5.**
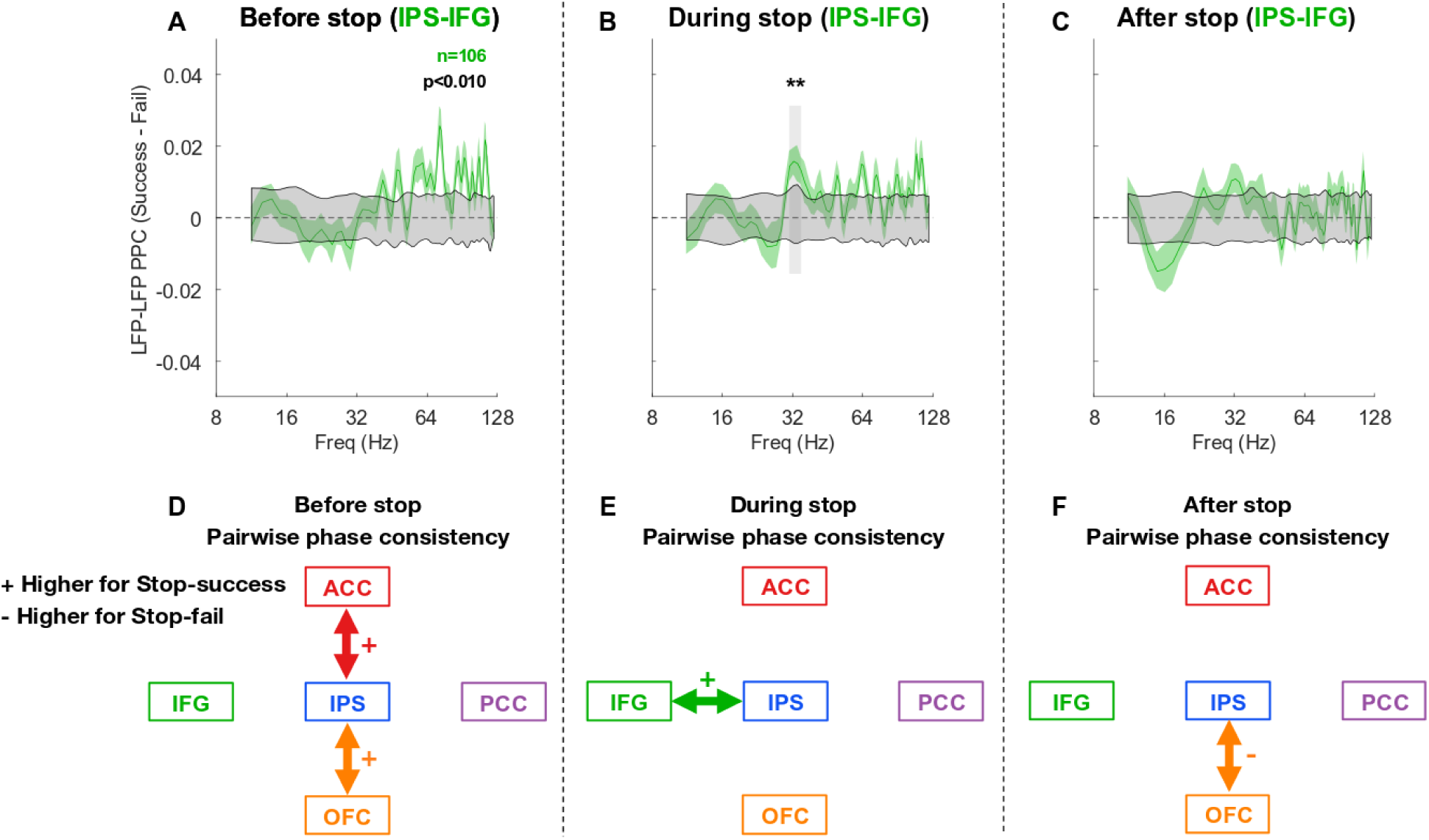
PPC between IPS and IFG in the SST across different time periods in the SST. We measured the difference in PPC between IPS and IFG in the (A) Before-stop, (B) During-stop, and (C) After-stop between Stop-success and Stop-fail trials. Positive values indicate that the PPC between IPS and IFG is higher during Stop-success than Stop-fail trials. (A-C) The black asterisks and gray vertical shaded region denote *p* < 0.01 (paired t-test); Gray shaded regions are the 99% bounds of a permutation test (see STAR Methods); Colored traces denote mean PPC values; Colored shaded regions denote ± SEM; n indicates the number of LFP-LFP pairs. (A) In the Before-stop epoch, there is no significant difference in PPC in successful and failed trials. (B) In the During-stop epoch, PPC between IPS and IFG is significantly higher in Stop-success trials at 31-35 Hz. (C) In the After-stop epoch, there is no significant PPC modulation. (D-F) A schematic of bidirectional information exchange between IPS and other brain regions based on LFP-LFP PPC. Double-sided arrow indicates significantly higher phase alignment in Stop-success (+ sign) and Stop-fail (- sign) trials, respectively. Details are shown in Figure S7. There is significant task-specific bidirectional communication in (D) IPS-OFC and IPS-ACC in the Before-stop epoch; (E) IPS-IFG in the During-stop epoch; and (F) IPS-OFC in the After-stop epoch, the only phase alignment effect that was higher for failed inhibition versus success.

Early in the trial, there was no significant difference in PPC values between IPS and IFG in Stop-success and Stop-fail trials (Figure 5A). However, in the During-stop epoch, there was significant modulation in PPC at 31-35 Hz (paired t-test; *p* < 0.01 at each frequency), which was not present anymore in the After-stop epoch (Figures 5B and 5C). We observed similar results when considering only the right hemisphere (Figure S6). We also quantified PPC between IPS and the other brain regions (Figures 5D-F and S7). There was significantly higher phase alignment for successful inhibition between IPS and OFC in the Before-stop epoch at 11-20 Hz (Figures 5D and S7A) and between IPS and ACC at 39-44 Hz (Figures 5D and S7C) (paired t-test; *p* < 0.01 at each frequency). The only effect associated with a higher phase alignment in Stop-fail trials was found between IPS and OFC in the After-stop epoch (44-48 Hz; Figures 5F and S7C).

## Discussion

We investigated the role of posterior parietal cortex in movement inhibition by recording intracranially in neurosurgical patients performing a standard SST. Inter-areal communication (e.g., functional or effective connectivity) is key to support complex cognitive functions,^34,94–97^ and dysfunctional connectivity has been increasingly linked to psychiatric conditions.^98–102^ Parietal cortex, considered the “crossroads” of the brain, could be a critical player and a potential neuromodulation target. In this study, we employed three different connectivity metrics to investigate the interactions between parietal cortex and other nodes important for inhibition, control and updating of actions, and internally-focused cognition. Our connectivity analyses help us unveil the complex dynamics between IPS and these areas in successful versus failed movement inhibition (Figure 6). Our functional connectivity results support prior evidence of a role of IPS in inhibition,^103–105^ and provide a novel insight into its function: altogether, IPS seems to be associated with action plan monitoring. IPS receives anterior inputs before stopping (from IFG, OFC and ACC), updates and sends feedback on correctly terminated actions to IFG, and receives input from OFC after mistakes. Directed connectivity from PCC, a DM node, is associated with lapses in control.

**Figure 6.**
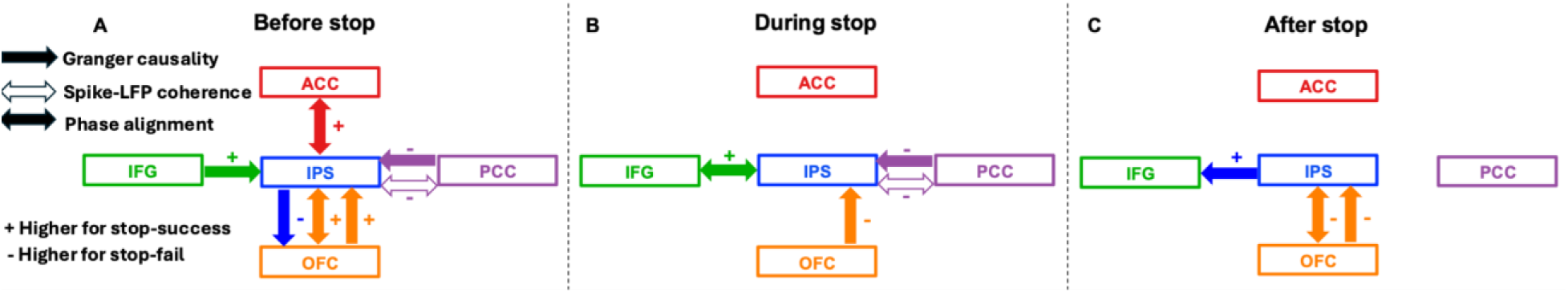
Summary schematic of communication between IPS and other brain regions. We depict a summary of all significant findings across the different metrics (different arrows) and denoting with “+” and “-” if the functional connectivity metric was higher for Stop-success or Stop-Fail. Note that connectivity results that were the same for success and fail are not included (e.g. ACC to IPS Granger causality results). (A) In the Before-stop epoch, IFG-to-IPS and OFC-to-IPS information flow is higher for Stop-success, and IPS-to-OFC and PCC-to-IPS information flow is higher for Stop-fail (Granger causality). Bi-directional communication between IPS and OFC and between ACC and IPS is higher for Stop-success (phase alignment). Higher PCC-to-IPS information flow in Granger causality for Stop-fail is consistent with spike-LFP coherence results. (B) In the During-stop epoch, OFC-to-IPS information flow is higher for Stop-fail (Granger causality). Bi-directional communication between IPS and IFG is higher for Stop-success (phase alignment). Communication between PCC and IPS is similar to the pattern in the Before-stop epoch in (A). (C) In the After-stop epoch, IPS-to-IFG information flow is higher for Stop-success, and OFC-to-IPS information flow is higher for Stop-fail (Granger causality). Bi-directional communication between IPS and OFC is higher for Stop-fail (phase alignment).

### IFG leads activity in parietal cortex, and their connectivity is critical to stopping

First, we confirmed the presence of differential activity between successful and failed stopping by comparing spectral power values across a wide time-frequency range for both posterior parietal cortex and IFG (Figure 1). Next, we used Granger causality to analyze patterns of information flow between IPS and IFG before, during and after the presentation of the stop signal. Successful movement inhibition was associated with IFG beta-range activity leading the communication with IPS when preparing to stop (Figure 2A). During stop signal presentation, phase alignment between the two areas was increased (Figure 5B). After that, the communication was reversed, with IPS leading IFG (Figure 2C). No notable communication asymmetry between IFG and IPS was detected by our Granger causality results during failures in control (see Figure 6 for a summary).

Previous work has proposed IFG as a starting point of the cascade of neural signals for inhibitory control forming a cortico-thalamic hyperdirect pathway.^106,107^ IPS has been understood as an area that does not come into play in the hyperdirect pathway, and instead participates in the inhibition of motor responses by ensuring accurate motor planning^18,108^ rather than reactive response inhibition.^17^ However, resting-state functional connectivity analyses between IFG and the IPS^37,109^ and tract tracing in the non-human primate brain^110,111^ showed evidence of direct connections between IFG and parietal locations. Further, recent fMRI and TMS studies on IPS during stop signal tasks indicated that there may be parallel processing streams in the cingulo-opercular and frontoparietal pathways for inhibitory control.^37,38^ Altogether, this evidence cannot rule out the possibility of IPS being a key node in the inhibition network. fMRI signals have low temporal resolution, making it impossible in a fast-paced paradigm such as SST to disentangle neuronal communication between brain regions.^112^ On the contrary, TMS has very precise timing, but poor spatial precision, and a conclusive consensus is lacking regarding the spatial extent of nervous tissue affected by TMS.^113,114^ Due to the spatial uncertainty together with variability in individual gyration, and excitability of cortical areas,^113,114^ one could wonder if TMS effects can be truly ascribed to IPS or if they are likely a compound effect from neighboring regions. However, Osada and colleagues showed an effect on inhibitory abilities (increased SSRT) for TMS interference on IPS, and not on the adjacent TPJ, thus demonstrating specificity.^38^ This was further ascribed to the presence of direct connections between IFG and IPS, but not IFG and TPJ.^38^ Our own findings provide further support for this interpretation. Indeed, we show functional connectivity during stopping between IFG and IPS, which is present only for successful trials. Thus, the disrupted inhibition performance caused by TMS over IPS could be explained by the interruption in the flow of information from frontal areas to IPS. These top-down influences from frontal areas to IPS demonstrate that parietal regions are recruited to aid inhibitory control,^45^ and that a lack of frontal-parietal communication is associated with lapses in control.^115^

Previous work used temporal latency and Granger causality analyses to infer information flow in movement inhibition between IFG and other areas. Timing of broadband gamma activity in intracranial signals was used to determine that IFG activity precedes that of anterior insula and of more anterior ventral prefrontal subdivisions.^73^ In our results, the latency of the beta band peak indicated that IFG activity preceded that of any other region. Schaum and colleagues measured information flow between IFG and pre-SMA based on magnetoencephalography signals,^80^ finding higher Granger causality from IFG to pre-SMA in movement inhibition. This result closely resembles the pattern we observed between IFG and IPS, with a key difference: there was no reversal in asymmetry in a later epoch from pre-SMA to IFG, while we found evidence of cross-talk from IPS to IFG. Overall, these asymmetric results from IFG to pre-SMA and from IFG to IPS support that IFG starts the cascade of neural signals that leads to stopping, and that IPS, like pre-SMA, is a recipient of that information. However, the fact that we observed increased communication from IPS to IFG in the After-stop epoch and that it was specific for successful inhibition demonstrates that IPS is more than just a passive receiver or executer. Given that IPS has the capability of influencing back IFG, it is possible that neuromodulation targeting IPS (i.e., TMS) would have cascading effects on the rest of the inhibitory control network. While it is difficult to speculate on the content of this communication, it is compatible with a feedback signal being relayed back to IFG from IPS. These could reflect the update of the action plan (interruption of the response to the Go cue, correct termination of the movement),^44^ but to support this interpretation recordings across the basal ganglia-primary motor pathway^74^ and IPS would be needed, disentangling the differential contributions of the frontoparietal and hyper-direct pathways in inhibition.

A unique feature of the SST is that with the staircase method we virtually obtain an equal number of success and error trials.^81^ When considering Stop-fail trials, we noted that the connectivity between IFG and IPS was not associated with any significant asymmetry. In opposition to this, we found complex patterns between IPS and the other recorded regions during control failures.

### Complex OFC and IPS connectivity changes might bias action plans and priorities

In Stop-success trials, there was higher information flow from OFC to IPS than vice versa in the Before-stop epoch in a low-beta band (Figures 3 and S3A). This higher communication between OFC and IPS was also shown in increased beta phase alignment (Figures 5 and S7A). In Stop-fail trials, however, information flows in the opposite direction in the Before-stop epoch, from IPS to OFC (Figures 3 and S4A). Thus, similar to what we observed for IFG, an early frontal to parietal asymmetry leads to successful inhibition. Later, during successful stopping, we find the opposite pattern, with OFC receiving more information from IPS (Figures 3 and S3B), while during stopping failures there is a mix of both directions (IPS to OFC in a high beta range, and OFC to IPS in low beta [Figures 3 and S4B]). Thus, this indicates functionally distinct channels of communication between these two areas, occupying separate spectral ranges within the beta band, with opposite effects on behavior depending on their timing: an early frontal to parietal directed connectivity associated with stopping, and an early parietal to OFC associated with failures in stopping (see Figure 6 for a summary). After-stop, no asymmetry is seen for successful stop, while there was an increase in OFC to IPS communication (Figures 3, S4B and S4C) paired with a higher phase alignment between the two regions for Stop-fail trials (Figures 5 and S7C). The role of OFC in inhibition has been classically attributed to its causal role in reversal learning, thus critical for inhibiting maladaptive and inappropriate responses^116^ and to map reward values on the appropriate responses.^67^ When considering our results on the early stages of stopping in this context, the OFC to IPS activity could reflect the OFC integrating prior with current information and communicating to IPS the optimal action-plan.^65,67^ A recent rodent study demonstrated a decrease in beta phase synchronization between OFC and STN in this stage, thus providing evidence of OFC influence over action maintenance and updating.^66^ Recently, the contributions of OFC to inhibition were reconciled with a broader decision making role,^67^ theorizing that OFC signals are not purely related to stopping, rather they bias behavior in a particular direction, untangling options^117^ and deciding^118,119^ which action to perform or prioritize. In our study, the during and post-error OFC to IPS activity could reflect the process of updating that information for better task performance on the next trial.^92,103,120^

### ACC connects to IPS to guide action selection

During both Stop-success and Stop-fail trials, there was higher information flow from ACC to IPS than vice versa (Figures 3, S3 and S4). This ACC to IPS communication was not only consistent regardless of trial outcome, but it was also consistent across different time windows, with some variations in the spectral range. The spike-LFP coherence analysis further confirmed the observation that the communication between these two regions did not depend on performance, as the alignment of ACC SUA with beta LFP in parietal cortex was not modulated by trial outcome (Figure 4). While this finding could seem at odds with the well-established role of ACC in monitoring and control,^121–124^ we find an indication of a modulation related to performance when considering the beta-band phase alignment, with higher synchronization between ACC to IPS for Stop-success in the window preceding the stop presentation (Figure 5). In addition, the influence of ACC over IPS is briefly increased following errors. Lastly, when considering the timing of the beta peak after the stop signal presentation, ACC was the only region to show a trend for an earlier beta-band peak for failed versus successful inhibition (Figure 3). Altogether, these results demonstrate that a brief phase alignment between ACC and IPS before the stop-signal is associated with correct inhibition (see Figure 6 for a summary), while the constant directional information flow from ACC to IPS is sustained regardless of performance, with a transient increase post-error. ACC is not considered a core region for inhibition, but more broadly important for interference resolution.^125^ The diverse roles played by this region across different cognitive functions have been proposed to underlie a general ability to track task-relevant information to guide appropriate actions.^126,127^ Our own results are consistent with this view: the sustained influence of ACC over IPS that seems to be independent of action outcomes, and only partially increased after an error, perhaps reflects the encoding of the task-state.^59,122,128^ The brief influence of ACC over IPS before stopping has behavioral consequences (i.e., interrupting the prepotent action), and the early latency of beta power following a stop-failure could be associated with task errors monitoring,^129^ in concert with the transient increase in connectivity between ACC and IPS.

### PCC to IPS communication is associated with lapses in control

Interactions between PCC and IPS were specific to stopping mistakes. Higher information flow from PCC to IPS was observed in both LFP-LFP Granger causality (Figures 3 and S4G) and spike-LFP coherence (Figure 4) in Stop-fail trials. PCC is typically considered part of the default mode network (DMN), a set of regions that is more active during resting state and internally-directed cognition.^130^ Our results indicate that higher activity from PCC to IPS is a detrimental feature, associated with lapses in control (see Figure 6 for a summary). A possible interpretation is that this represents a shift in the focus of attention,^131^ moving from the task stimuli and cues toward internal states (DM-related), effectively disengaging the IPS (and perhaps more broadly the frontoparietal network) from the ongoing performance and its monitoring.^69,132,133^ There is an increasing amount of evidence that complex cognitive functions rely on the interplay and switching of activity across multiple networks, including the DMN.^134,135^ For example, previous work from our group demonstrated that DMN plays a causal role in creative thinking, ^136,137^ and that in the early stages of this complex ability there is a simultaneous recruitment of the frontoparietal and DM networks. From this perspective, the cross-network interactions reported here may represent a suboptimal state,^138^ with PCC’s influence over IPS interfering with the task-relevant interactions with IFG, ACC and OFC and causing lapses in control^138^ leading to errors.^131^ An alternative interpretation is related to the evidence of an executive role of PCC,^139^ potentially spatially segregated from the PCC DM region,^71^ engaged by risky choices^140^ and prediction errors.^141^ With the current dataset we cannot discriminate if the negative influence of PCC is the cause for failures (internally-directed cognition interferences, causing the participant to fail at stopping) or if it represents a correlate of encoding prediction errors, needed to alter future actions, in line with a more executive role. Nonetheless, we provide novel evidence of an interaction between posterior cingulate and parietal cortex, evident also at the neuronal level, associated with failures in inhibitory control.

### Conclusions and Future Directions

In the current study, we showed performance-specific communication between IPS and other brain regions. The rich patterns of connections to and from IPS could indicate the potential of this area to be considered as a new therapeutic target for patients with inhibitory control dysfunctions. Previous work showed that neuromodulation of parietal cortex^142^ primarily induced psychophysical effects such as motor intention,^52,143^ visuomotor integration,^144^ attention,^145–148^ memory,^149,150^ and improved psychiatric symptoms of anxiety and depression.^151–153^ In particular, Balderston and colleagues used IPS as their therapeutic target to reduce physiological arousal in anxiety^153^ motivated by their previous findings of a higher global IPS connectivity in threat or shock.^154^ Our work expands from previous evidence of a causal role of IPS on stopping abilities,^37,38^ and reveals its complex pattern of functional interconnections. Through IPS neuromodulation, we may be able to tap into the ability of monitoring and updating action plans, with the advantage of a tractable and accessible target for non-invasive techniques.

### Limitations of the study

The primary focus of the current study was on LFP-LFP and spike-LFP interactions. Yet, Granger causality measures, despite its nomenclature, are not truly causal metrics. Stimulation methods are necessary to establish causality and give us further insight into the functional roles of inter-areal communication.^155^ While our work is in agreement with the few existing studies leveraging TMS and tDCS to assess the functional contribution of parietal regions in inhibitory control,^37,38,45^ more research is needed to disentangle the relative and causal contributions of inter-areal communication to inhibitory control. Indeed, it is difficult to separate what is truly “inhibition specific” from domain-general functions that play a role in inhibition, and further causal manipulations will be needed to provide us with more conclusive evidence about the role of IPS and its potential as a neuromodulation target for impulse control disorders.

Our inter-areal analyses all focus on interactions to and from parietal cortex, not testing direct interactions between other regions (i.e., not mediated by parietal signals). While those interactions are beyond the scope of the current study, more studies with temporally-resolved interactions between multiple regions are needed to further disentangle the flow of activity underlying inhibitory control.^66,73,74,80^

## STAR METHODS

### KEY RESOURCES TABLE

| REAGENT or RESOURCE | SOURCE | IDENTIFIER |
| --- | --- | --- |
| **Deposited Data** |  |  |
| Single unit, LFP, LFP-LFP Granger causality, LFP-LFP PPC, Spike-LFP coherence between brain regions | Current study | TBA |
| **Experiment models: Organisms/strains** |  |  |
| Human subjects |  |  |
| **Software and algorithms** |  |  |
| Fieldtrip | Fieldtrip Developers | https://www.fieldtriptoolbox.org/ |
| Freesurfer v6.0 | Freesurfer Developers | http://freesurfer.net/ |
| FSL | FSL Developers | https://fsl.fmrib.ox.ac.uk/fsl/docs/#/ |
| iELVis | iELVis Developers | http://ielvis.pbworks.com/w/page/116347253/FrontPage |
| MATLAB | Mathworks | https://www.mathworks.com/ |
| Psychtoolbox-3 functions (v3.0.16) | Psychtoolbox Developers | http://psychtoolbox.org/ |
| R | The R Foundation for Statistical Computing | https://www.r-project.org/ |
| Original code for data analysis | Current study | TBA |
| **Other** |  |  |
| sEEG probes | PMT, MN, USA | http://www.pmtcorp.com/ |
| BlackRock Neuroport system | BlackRock Microsystems, UT, USA | https://blackrockmicro.com/ |

### RESOURCE AVAILABILITY

#### Lead Contact

Requests for further information should be directed to the lead contact, Eleonora Bartoli.

#### Materials availability

N/A. No reagents were generated in this study.

#### Data and code availability

- All data reported in this paper will be shared by the lead contact upon request.
- All original code will be deposited at Zenodo and will be publicly available at the date of publication.
- Any additional information required to reanalyze the data reported in this paper is available from the lead contact upon request.

### EXPERIMENTAL MODEL AND SUBJECT DETAILS

12 subjects (6 males, 6 females, mean age of 26 years, ranging from 24-53 years) consented to participate in this study while they underwent invasive epilepsy monitoring with sEEG at Baylor St. Luke’s Medical Center (Houston, TX, USA). Detailed subject information is reported in Table S1. Experimental procedures approved by the Institutional Review Board at Baylor College of Medicine (IRB protocol number H-18112). No patients with prior surgical resection in the areas of interest participated in this study. Experiments were recorded while interictal epileptic discharges were minimal and no seizures occurred within 2 hours of the experiments.

### METHOD DETAILS

#### Task design

All participants performed the experimental paradigm while reclined in a hospital bed in a quiet room. All tasks were presented on an adjustable monitor (1920×1080 resolution, 47.5×26.7 cm screen size, 60Hz refresh rate, connected to a PC running Windows 10 Pro at a viewing distance of 57 cm, such that 1 cm on the screen corresponds to ∼1 degree visual angle). Tasks were programmed using Psychtoolbox-3 functions (v3.0.16)^156^ running on MATLAB (R2019a, MathWorks, MA, USA).

Participants performed a simple stop signal task paradigm, adapted from the “STOP-IT2_beta03” software.^82^ Each trial started with a fixation cross (500 ms duration) followed by a go cue (an arrow pointing to the left or to the right, 1,500 ms duration). The participant was instructed to press the corresponding button on a keyboard (with the right hand). On 25% of trials, a stop signal (color change in the arrow, from gray to bright orange) was presented at a variable delay from the onset of the go cue (stop signal delay, SSD) and participants attempted to stop the button press response. Each participant completed a brief training session (10 trials with feedback) to familiarize with the task and receive coaching. This data was not analyzed. The experimental session consisted of 3 blocks of 96 trials each separated by short breaks (10 seconds), with the stop signal being presented one fourth of the trials (of 288 total trials, 216 go and 72 stop trials). The SSD was initially set at 250 ms and varied adaptively based on performance with the staircase method (SSD increased by 50 ms with each successful stop, decreased by 50 ms after a failed stop, with SSD boundaries set at 0 and 1,300 ms). Two independent staircases were used for the left and right button press responses. Participants repeated the whole experimental session a second time with a different visual background. Here we report data from only one session, as the visual background manipulation is not relevant to the current study.

#### Behavioral analysis

For each trial, the following measures were collected: accuracy and, if a response was given, reaction time. From these metrics we computed: the probability of responding to a stop signal (number of “Stop-fail” trials over number of stop trials), accuracy on Go trials, reaction time during Go trials, and reaction time in “Stop-fail” trials. We ensured that the assumptions of the race model were met by testing that the reaction times for failed inhibition were shorter than the ones for go trials using a non-parametric paired test (Wilcoxon signed rank test). The stop signal reaction time (SSRT) was estimated using the integration method.

#### Stereoelectroencephalography (sEEG) probes

Reported data were acquired by sEEG depth probes. The sEEG probes had either a 0.8 mm diameter, with 8 to 16 electrode contacts along the probe with a 3.5 mm center-to-center distance (PMT Corporation, MN, USA) or a 1.28 mm diameter, with 9 recording contacts, with a 5.0 mm center-to-center distance between contacts (AdTech Medical Instrument Corporation, WI, USA).

#### Electrode localization and selection

For each patient, we determined electrode locations by employing the software pipeline intracranial Electrode Visualization, iELVis.^157^ In short, the post-operative computed tomography (CT) image was registered to the pre-operative T1 anatomical magnetic resonance imaging (MRI) image using FSL.^158^ Next, the location of each electrode was identified in the CT-MRI overlay using BioImage Suite.^159^ Anatomical locations of the electrodes were classified based on their proximity to the cortical surface model, reconstructed by the T1 image using Freesurfer (version 6.0).^160^

We further labeled each electrode by finding the most likely cortical parcellation estimate in a 5 mm radius around each electrode using the Destrieux Cortical Atlas.^161^ The atlas contains 76 cortical parcellation labels and each cortical surface point is assigned a label by mapping the parcellation estimates on each individual cortical surface. The inclusion criteria for this study was the presence of at least one electrode recording from posterior parietal cortex (Destrieux labels: ‘G_pariet_inf-Angular’, ‘G_pariet_inf-Supramar’, ‘G_pariet_sup’, ‘G_precuneus’, ‘S_interm_prim-Jensen’, ‘S_intrapariet_and_P_trans’, ‘S_parieto_occipital’, ‘S_subparietal’). Electrodes classified as recording from the IFG (Destrieux labels containing ‘inf-Triangul’), IPS (‘intrapariet’), OFC (‘orbital’), ACC (‘cingul-Mid-Ant’ and ‘cingul-Ant’), and PCC (‘S_cingul-Marginali’, ‘S_subparietal’, ‘S_parieto_occipital’) were selected for the main functional connectivity analyses of the current study.^162^ For the spike-LFP coherence analysis, we selected SUA isolated from microwires recording from ACC and PCC (defined above) and LFP recorded from parietal cortex (‘pariet’, ‘Jensen’, ‘Marginalis’, ‘precuneus’).

#### Electrophysiological recording and preprocessing

sEEG signals were recorded by a 256 channel BlackRock Neuroport system (BlackRock Microsystems, UT, USA) at 2 kHz sampling rate, with a 4^th^ order Butterworth bandpass filter of 0.3-500 Hz. sEEG recordings were referenced to an electrode contact visually determined to be in white matter and at least 3 mm away from gray. A photoresistor sensor recording at 30 kHz was attached to the task monitor to synchronize intracranial recordings to the precise go and stop signal presentation time (a white square was presented at the same time as the go and stop cues in the lower right corner of the monitor, and its timing was recorded through the photoresistor signal). All signals were processed in MATLAB (R2019a, MathWorks, MA, USA). Raw sEEG signals were firstly inspected for line noise, recording artifacts (e.g., contamination by muscle activity, presence of large deflections, additional noise components), and interictal epileptic spikes. Electrodes with artifacts and epileptic spikes were excluded from further analysis. Next, the signal from each electrode was re-referenced to the average of all electrodes that survived exclusion criteria. Re-referenced signals from electrodes localized within our ROIs were selected for further analysis. 12 patients had parietal coverage and were included in the study. Across this sample, 7 participants had electrodes recording IPS and 6 from both IPS and IFG. Table S2 has detailed information about the number of electrodes in brain regions of interest for each patient.

Single-unit activity (SUA) was obtained in a subset of locations by Behnke-Fried configuration hybrid macro-micro electrodes, with a bundle of 8 recording shielded microwires at the tip of the electrode (∼40 μm diameter, extending about 3 mm from the tip) and 1 unshielded microwire used as a local reference. Signals from Behnke-Fried electrodes were recorded at 30 kHz using NeuroSnap Headstages connected to a NeuroSnap Headstage Bridge, feeding into the BlackRock Neuroport system. Across our sample, 6 participants had microwires in ACC and at least one sEEG electrode in parietal, and 3 of those also had microwires in PCC. In this work, we refer to LFP as the signal recorded from macro-electrodes (sEEG) only. From the micro-electrodes, only SUA was analyzed, as described below in spike-LFP PPC.

#### Electrophysiological analysis

Re-referenced signals were downsampled to 1000 Hz and spectrally decomposed using a family of Morlet wavelets (7 cycles), with center frequencies ranging logarithmically from 1 to 200 Hz in 100 steps. First, we performed a time-frequency analysis of LFP power from all locations recording from the posterior parietal cortex and from IFG, and compared power values associated with Stop-success and Stop-fail. We identified the time of the stop signal presentation (using the photoresistor signal) and we selected time-frequency power values from the 300 ms to 700 ms after the stop sign. Power values during each stop trial were normalized to percent change with respect to the pre-trial baseline period (300 to 0 ms before go cue presentation). Power % change values computed for each electrode location were then averaged over Stop-success trials and Stop-fail trials separately, for each electrode location and participant. To assess the presence of differences in the time-frequency power values, we employed permutation testing (5,000 permutations), randomly shuffling labels between Stop-success and Stop-fail to estimate the distribution of differences at each time-frequency point under the null hypothesis and compute p-values. More details in the statistics section below. For LFP latency detection, we focused on the beta range. Beta band power was obtained by averaging the magnitude of the Morlet wavelet decomposition result between 15-25 Hz for beta, and normalized to percent change with respect to the base-line period (300 to 0 ms before stop cue presentation) (Figure S8). Next, for each electrode location, we used the matlab function *findpeaks* in MATLAB and we defined the first beta peak as the first local maxima after stop cue onset. We adjusted the parameter *MinPeakProminence* to 10 to filter out minor variation and noise. We used this setting after visually inspecting LFP power traces for reliable peak detection across all LFP data.

#### Spectral LFP-LFP Granger causality

We recorded LFP-LFP pairs between IPS and IFG (n=106 pairs; 6 patients), IPS and OFC (n=247 pairs; 5 patients), IPS and ACC (n=231; 7 patients), and IPS and PCC (n=233; 4 patients). To quantify causal relationships between LFP signals in different brain regions, we implemented spectral Granger causality (nonparametric bivariate) using the FieldTrip toolbox.^163^ Granger causality in neural signals estimates causal influence from signal *X* to signal *Y* by comparing the variance of signal *Y* that can be explained by the recent history of the signal *Y* alone with the variance that can be explained by taking the recent history of both signals *X* and *Y* into account.^164^ If the difference between the two variances is large, we interpret that there is causal influence from signal *X* to signal *Y*. Time-based autoregressive models, however, may not fully reflect spectral qualities of signals.^165,166^ Because we aimed to study frequency-specific causal influences between LFP signals, we implemented spectral LFP-LFP Granger causality from signal *X* to signal *Y* at frequency *ω* to quantify casual relationships using cross-spectral density matrix factorization in Eqs. (1) and (2).^83,167^

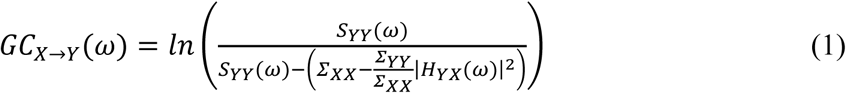

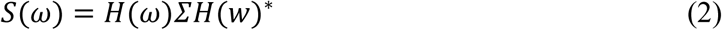

For a pair of signals *X* and *Y* at frequency *ω*, *S*(*ω*) is the cross-spectral density matrix and *H*(*ω*) is the spectral transfer matrix. *Σ* is the covariance of an autoregressive model’s residuals. We excluded LFP signals below 10 Hz because they were unreliable (Figure S1).^83,91^

#### LFP-LFP pairwise-phase consistency (PPC)

To measure synchronization of two LFP signals, we implemented LFP-LFP PPC^93^ using the FieldTrip toolbox.^163^ Compared to phase coherence, PPC is less affected by amplitude correlations and therefore is a bias-free metric for LFP synchronization.^93,168^ PPC of LFP signals *X* and *Y* at frequency *ω* was estimated as in Eq. (3).

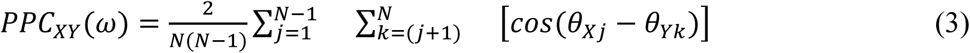

*N* is a total number of cycles at frequency *ω*. *θ*_*Xj*_ and *θ*_*Yk*_ are unit vectors that correspond to phase angles of signals *X* and *Y* at a particular cycle of *j* and *k*, respectively.

#### Spike-LFP PPC

To measure synchronization of spikes and LFP signals, we implemented spike-LFP PPC using the FieldTrip toolbox.^163^ Spike-LFP PPC is a bias-free statistic that is not affected by the number of spikes.^88,93^ We measured consistency of phases of from all LFP signals 100 ms before and 100 ms after each spike (200 ms sliding window with a stepping size of 10 ms) in 15, 20, and 25 Hz. For this analysis, we used spikes detected during the whole trial [-500 ms +1500 ms] aligned to the onset of the first black arrow. We chose a broader epoch to ensure higher spike counts and robust statistics. Spike-LFP PPC at frequency *ω* of between spikes and and a LFP signal was estimated as in Eq. (4)

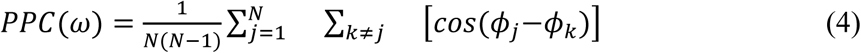

*N* is a total number of spikes. *ϕ*_*j*_ and *ϕ*_*k*_ are the phases of the LFP signal in the 200 ms time window at the time of the *j*-th and *k*-th spike, respectively. Extracellular spikes were detected on the band-passed microwire signals (300-3000 Hz) based on a signal-dependent negative voltage threshold, and automatically sorted employing waveform features and unsupervised nonparametric clustering (Wave_clus3).^169^ Isolated waveforms were segregated into units in a semi-automated manner. The times of threshold crossing for identified units were retained for spike-LFP coherence analysis. Based on previously used criterion,^170^ units were classified as single- or multi-units based on the spike shape and inter-spike-interval (ISI) distribution of the cluster (single-units with <1% of spikes with an ISI less than 3 ms). In addition, to be included in the spike-LFP PPC analysis, units needed to have a sufficient number of spikes across the stop trials (minimum n = 72). Across the 6 subjects with Behnke-Fried probes in ACC, 62 single-units were used for this analysis following the inclusion criteria. From the 3 subjects with probes in PCC, 12 single-units were employed. The number of spike-LFP pairs between participants ranged from 8-1036 (ACC- parietal cortex) and 12 to 40 (PCC-parietal cortex).

#### Statistics

Statistical analyses were conducted in MATLAB (Mathworks) and R Statistical Software (4.3.3). All statistical tests were two-sided unless specified as one-sided. Statistical significance for time-frequency power values differences was obtained using permutation testing (5,000 permutations) and using cluster-based correction for multiple comparisons across the time-frequency space. Clusters associated with a corrected *p* < 0.05 were considered significant. To test for statistical significance between behavioral measures and connectivity metrics (Figures 2, 3, 4, 5, S1, S2, S3, S4, S6, S7), we applied either Wilcoxon signed-rank or paired t-tests. Granger causality is a biased metric with a minimum of 0 but no upper bound for maximum, and we used Wilcoxon signed-rank tests for statistics of Granger causality values. Multiple comparisons correction for spectral measures is challenging because values at neighboring frequencies are not independent. We also have a step where we do frequency smoothing with a Gaussian filter in our analysis, and this step increases dependencies between adjacent frequencies. Therefore, we used a conservative statistical threshold for our statistical testing: *p* < 0.01 in four or more contiguous bands (below 60 Hz) and six or more contiguous bands (above 60 Hz). For the latency analysis, we employed a mixed effect model approach (*lmer* package in R) defining fixed effects for electrode location (levels: IPS, IFG, ACC, OFC, PCC), and trial type (Success and Failure) and random effects for participants. P-values were approximated using Satterthwaite’s method. Post-hoc comparisons were used to test for differences between Stop-success and Stop-fail trials within each region (*emmeans* function *contrast* with Bonferroni correction for multiple comparisons). Statistical significance is denoted as \**p* < 0.05 and \*\**p* < 0.01.

## Acknowledgements

This work was supported by the Robert and Janice McNair Foundation (S.A.S.) and by the NIH (UH3NS103549 and R01MH130597). J.V. was supported by the Secretaría de Educación, Ciencia, Tecnología e Innovación de la Ciudad de México (Grant Number: SECTEI/103/2022).

## Author contributions

Conceptualization: E.B. Data collection: L.M., J.V., H.G.R., A.J.W., and E.B. Data analysis: J.K., V.E.G., and E.B. Original manuscript: J.K. Feedback for manuscript revision: All authors. Project supervision: E.B., S.A.S., and B.Y.H. Funding acquisition: S.A.S.

## References

1. Botvinick, M.M., Braver, T.S., Barch, D.M., Carter, C.S., and Cohen, J.D. (2001). Conflict monitoring and cognitive control. Psychol. Rev. 108, 624–652. 10.1037/0033-295X.108.3.624.

2. Braver, T.S. (2012). The variable nature of cognitive control: a dual mechanisms framework. Trends Cogn. Sci. 16, 106–113. 10.1016/j.tics.2011.12.010.

3. Miller, E.K. (2000). THE PREFRONTAL CORTEX AND COGNITIVE CONTROL. Nat. Rev. Neurosci. 1, 59–65. 10.1038/35036228.

4. Logan, G.D. (1985). Executive control of thought and action. Acta Psychol. (Amst.) 60, 193–210. 10.1016/0001-6918(85)90055-1.

5. Aron, A.R. (2007). The Neural Basis of Inhibition in Cognitive Control. The Neuroscientist 13, 214–228. 10.1177/1073858407299288.

6. Wessel, J.R., and Anderson, M.C. (2024). Neural mechanisms of domain-general inhibitory control. Trends Cogn. Sci. 28, 124–143. 10.1016/j.tics.2023.09.008.

7. Logan, G.D., and Cowan, W.B. (1984). On the ability to inhibit thought and action: A theory of an act of control. Psychol. Rev. 91, 295–327. 10.1037/0033-295X.91.3.295.

8. Wessel, J. (2023). Action stopping. Preprint at OSF, 10.31234/osf.io/9j63e https://doi.org/10.31234/osf.io/9j63e.

9. Lustig, C., Hasher, L., and Tonev, S.T. (2001). Inhibitory control over the present and the past. Eur. J. Cogn. Psychol. 13, 107–122. 10.1080/09541440126215.

10. Pearce, B., Cartwright, J., Cocks, N., and Whitworth, A. (2016). Inhibitory control and traumatic brain injury: The association between executive control processes and social communication deficits. Brain Inj. 30, 1708–1717. 10.1080/02699052.2016.1202450.

11. Zhao, L., Silva, A.B., Kurteff, G.L., and Chang, E.F. (2023). Inhibitory control of speech production in the human premotor frontal cortex. Preprint, 10.1101/2023.03.01.530698 https://doi.org/10.1101/2023.03.01.530698.

12. Zheng, X., Roelofs, A., Erkan, H., and Lemhöfer, K. (2020). Dynamics of inhibitory control during bilingual speech production: An electrophysiological study. Neuropsychologia 140, 107387. 10.1016/j.neuropsychologia.2020.107387.

13. Ganos, C., Rothwell, J., and Haggard, P. (2018). Voluntary inhibitory motor control over involuntary tic movements. Mov. Disord. 33, 937–946. 10.1002/mds.27346.

14. Jurgiel, J., Miyakoshi, M., Dillon, A., Piacentini, J., Makeig, S., and Loo, S.K. (2021). Inhibitory control in children with tic disorder: aberrant fronto-parietal network activity and connectivity. Brain Commun. 3, fcab067. 10.1093/braincomms/fcab067.

15. Mancini, C., Cardona, F., Baglioni, V., Panunzi, S., Pantano, P., Suppa, A., and Mirabella, G. (2018). Inhibition is impaired in children with obsessive-compulsive symptoms but not in those with tics. Mov. Disord. 33, 950–959. 10.1002/mds.27406.

16. Aron, A.R. (2011). From Reactive to Proactive and Selective Control: Developing a Richer Model for Stopping Inappropriate Responses. Biol. Psychiatry 69, e55–e68. 10.1016/j.biopsych.2010.07.024.

17. Wessel, J.R., and Aron, A.R. (2017). On the Globality of Motor Suppression: Unexpected Events and Their Influence on Behavior and Cognition. Neuron 93, 259–280. 10.1016/j.neuron.2016.12.013.

18. Hannah, R., and Aron, A.R. (2021). Towards real-world generalizability of a circuit for action-stopping. Nat. Rev. Neurosci. 22, 538–552. 10.1038/s41583-021-00485-1.

19. Wyland, C.L., Kelley, W.M., Macrae, C.N., Gordon, H.L., and Heatherton, T.F. (2003). Neural correlates of thought suppression. Neuropsychologia 41, 1863–1867. 10.1016/j.neuropsychologia.2003.08.001.

20. Li, C.R., Huang, C., Yan, P., Bhagwagar, Z., Milivojevic, V., and Sinha, R. (2008). Neural Correlates of Impulse Control During Stop Signal Inhibition in Cocaine-Dependent Men. Neuropsychopharmacology 33, 1798–1806. 10.1038/sj.npp.1301568.

21. Versace, F., Robinson, J.D., and Cinciripini, P.M. (2023). Toward neuromarkers for tailored smoking cessation treatments. Addict. Neurosci. 6, 100075. 10.1016/j.addicn.2023.100075.

22. Luigjes, J., Segrave, R., de Joode, N., Figee, M., and Denys, D. (2019). Efficacy of Invasive and Non-Invasive Brain Modulation Interventions for Addiction. Neuropsychol. Rev. 29, 116–138. 10.1007/s11065-018-9393-5.

23. Spagnolo, P.A., and Goldman, D. (2017). Neuromodulation interventions for addictive disorders: challenges, promise, and roadmap for future research. Brain 140, 1183–1203. 10.1093/brain/aww284.

24. Kelley, N.J., Gallucci, A., Riva, P., Romero Lauro, L.J., and Schmeichel, B.J. (2019). Stimulating Self-Regulation: A Review of Non-invasive Brain Stimulation Studies of Goal-Directed Behavior. Front. Behav. Neurosci. 12. 10.3389/fnbeh.2018.00337.

25. Verbruggen, F., Aron, A.R., Band, G.P., Beste, C., Bissett, P.G., Brockett, A.T., Brown, J.W., Chamberlain, S.R., Chambers, C.D., Colonius, H., et al. (2019). A consensus guide to capturing the ability to inhibit actions and impulsive behaviors in the stop-signal task. eLife 8, e46323. 10.7554/eLife.46323.

26. Schall, J.D. (2001). Neural basis of deciding, choosing and acting. Nat. Rev. Neurosci. 2, 33–42. 10.1038/35049054.

27. Eijsker, N., Schröder, A., Smit, D.J.A., Van Wingen, G., and Denys, D. (2019). Neural Basis of Response Bias on the Stop Signal Task in Misophonia. Front. Psychiatry 10, 765. 10.3389/fpsyt.2019.00765.

28. Schall, J.D., Stuphorn, V., and Brown, J.W. (2002). Monitoring and Control of Action by the Frontal Lobes. Neuron 36, 309–322. 10.1016/S0896-6273(02)00964-9.

29. Swann, N., Tandon, N., Canolty, R., Ellmore, T.M., McEvoy, L.K., Dreyer, S., DiSano, M., and Aron, A.R. (2009). Intracranial EEG Reveals a Time- and Frequency-Specific Role for the Right Inferior Frontal Gyrus and Primary Motor Cortex in Stopping Initiated Responses. J. Neurosci. 29, 12675–12685. 10.1523/JNEUROSCI.3359-09.2009.

30. Aron, A.R., Robbins, T.W., and Poldrack, R.A. (2004). Inhibition and the right inferior frontal cortex. Trends Cogn. Sci. 8, 170–177. 10.1016/j.tics.2004.02.010.

31. Aron, A.R., Robbins, T.W., and Poldrack, R.A. (2014). Inhibition and the right inferior frontal cortex: one decade on. Trends Cogn. Sci. 18, 177–185. 10.1016/j.tics.2013.12.003.

32. Swann, N.C., Cai, W., Conner, C.R., Pieters, T.A., Claffey, M.P., George, J.S., Aron, A.R., and Tandon, N. (2012). Roles for the pre-supplementary motor area and the right inferior frontal gyrus in stopping action: Electrophysiological responses and functional and structural connectivity. NeuroImage 59, 2860–2870. 10.1016/j.neuroimage.2011.09.049.

33. Zhang, R., Geng, X., and Lee, T.M.C. (2017). Large-scale functional neural network correlates of response inhibition: an fMRI meta-analysis. Brain Struct. Funct. 222, 3973– 3990. 10.1007/s00429-017-1443-x.

34. Cole, M.W., Yarkoni, T., Repovs, G., Anticevic, A., and Braver, T.S. (2012). Global Connectivity of Prefrontal Cortex Predicts Cognitive Control and Intelligence. J. Neurosci. 32, 8988–8999. 10.1523/JNEUROSCI.0536-12.2012.

35. Harding, I.H., Yücel, M., Harrison, B.J., Pantelis, C., and Breakspear, M. (2015). Effective connectivity within the frontoparietal control network differentiates cognitive control and working memory. NeuroImage 106, 144–153. 10.1016/j.neuroimage.2014.11.039.

36. Scolari, M., Seidl-Rathkopf, K.N., and Kastner, S. (2015). Functions of the human frontoparietal attention network: Evidence from neuroimaging. Curr. Opin. Behav. Sci. 1, 32–39. 10.1016/j.cobeha.2014.08.003.

37. Osada, T., Ogawa, A., Suda, A., Nakajima, K., Tanaka, M., Oka, S., Kamagata, K., Aoki, S., Oshima, Y., Tanaka, S., et al. (2021). Parallel cognitive processing streams in human prefrontal cortex: Parsing areal-level brain network for response inhibition. Cell Rep. 36, 109732. 10.1016/j.celrep.2021.109732.

38. Osada, T., Ohta, S., Ogawa, A., Tanaka, M., Suda, A., Kamagata, K., Hori, M., Aoki, S., Shimo, Y., Hattori, N., et al. (2019). An Essential Role of the Intraparietal Sulcus in Response Inhibition Predicted by Parcellation-Based Network. J. Neurosci. 39, 2509–2521. 10.1523/JNEUROSCI.2244-18.2019.

39. Garavan, H., Ross, T.J., and Stein, E.A. (1999). Right hemispheric dominance of inhibitory control: An event-related functional MRI study. Proc. Natl. Acad. Sci. 96, 8301– 8306. 10.1073/pnas.96.14.8301.

40. Dodds, C.M., Morein-Zamir, S., and Robbins, T.W. (2011). Dissociating Inhibition, Attention, and Response Control in the Frontoparietal Network Using Functional Magnetic Resonance Imaging. Cereb. Cortex 21, 1155–1165. 10.1093/cercor/bhq187.

41. Aron, A.R., and Poldrack, R.A. (2006). Cortical and Subcortical Contributions to Stop Signal Response Inhibition: Role of the Subthalamic Nucleus. J. Neurosci. 26, 2424–2433. 10.1523/JNEUROSCI.4682-05.2006.

42. Gavazzi, G., Giovannelli, F., Currò, T., Mascalchi, M., and Viggiano, M.P. (2021). Contiguity of proactive and reactive inhibitory brain areas: a cognitive model based on ALE meta-analyses. Brain Imaging Behav. 15, 2199–2214. 10.1007/s11682-020-00369-5.

43. Gavazzi, G., Giovannelli, F., Noferini, C., Cincotta, M., Cavaliere, C., Salvatore, M., Mascalchi, M., and Viggiano, M.P. (2023). Subregional prefrontal cortex recruitment as a function of inhibitory demand: an fMRI metanalysis. Neurosci. Biobehav. Rev. 152, 105285. 10.1016/j.neubiorev.2023.105285.

44. Hannah, R., and Jana, S. (2019). Disentangling the role of posterior parietal cortex in response inhibition. J. Neurosci. 39, 6814–6816. 10.1523/JNEUROSCI.0785-19.2019.

45. Shiga, K., Miyaguchi, S., Inukai, Y., Otsuru, N., and Onishi, H. (2023). Transcranial direct current stimulation over the right intraparietal sulcus improves response inhibition. Behav. Brain Res. 437, 114110. 10.1016/j.bbr.2022.114110.

46. Vivas, A.B., Humphreys, G.W., and Fuentes, L.J. (2003). Inhibitory processing following damage to the parietal lobe. Neuropsychologia 41, 1531–1540. 10.1016/S0028-3932(03)00063-0.

47. Colby, C.L., and Goldberg, M.E. (1999). SPACE AND ATTENTION IN PARIETAL CORTEX. Annu. Rev. Neurosci. 22, 319–349. 10.1146/annurev.neuro.22.1.319.

48. Behrmann, M., Geng, J.J., and Shomstein, S. (2004). Parietal cortex and attention. Curr. Opin. Neurobiol. 14, 212–217. 10.1016/j.conb.2004.03.012.

49. Corbetta, M., and Shulman, G.L. (2002). Control of goal-directed and stimulus-driven attention in the brain. Nat. Rev. Neurosci. 3, 201–215. 10.1038/nrn755.

50. Capotosto, P., Corbetta, M., Romani, G.L., and Babiloni, C. (2012). Electrophysiological Correlates of Stimulus-driven Reorienting Deficits after Interference with Right Parietal Cortex during a Spatial Attention Task: A TMS-EEG Study. J. Cogn. Neurosci. 24, 2363– 2371. 10.1162/jocn_a_00287.

51. Sengupta, A., Banerjee, S., Ganesh, S., Grover, S., and Sridharan, D. (2024). The right posterior parietal cortex mediates spatial reorienting of attentional choice bias. Nat. Commun. 15, 6938. 10.1038/s41467-024-51283-z.

52. Desmurget, M., Epstein, C.M., Turner, R.S., Prablanc, C., Alexander, G.E., and Grafton, S.T. (1999). Role of the posterior parietal cortex in updating reaching movements to a visual target. Nat. Neurosci. 2, 563–567. 10.1038/9219.

53. Gréa, H. (2002). A lesion of the posterior parietal cortex disrupts on-line adjustments during aiming movements. Neuropsychologia 40, 2471–2480. 10.1016/S0028-3932(02)00009-X.

54. Scott, S.H. (2004). Optimal feedback control and the neural basis of volitional motor control. Nat. Rev. Neurosci. 5, 532–545. 10.1038/nrn1427.

55. Jana, S., Hannah, R., Muralidharan, V., and Aron, A.R. (2020). Temporal cascade of frontal, motor and muscle processes underlying human action-stopping. eLife 9, e50371. 10.7554/eLife.50371.

56. Hens, C., Harush, U., Haber, S., Cohen, R., and Barzel, B. (2019). Spatiotemporal signal propagation in complex networks. Nat. Phys. 15, 403–412. 10.1038/s41567-018-0409-0.

57. Jayasinghe, S.A.L., Scheidt, R.A., and Sainburg, R.L. (2022). Neural Control of Stopping and Stabilizing the Arm. Front. Integr. Neurosci. 16. 10.3389/fnint.2022.835852.

58. Whelan, R., Conrod, P.J., Poline, J.-B., Lourdusamy, A., Banaschewski, T., Barker, G.J., Bellgrove, M.A., Büchel, C., Byrne, M., Cummins, T.D.R., et al. (2012). Adolescent impulsivity phenotypes characterized by distinct brain networks. Nat. Neurosci. 15, 920–925. 10.1038/nn.3092.

59. Braver, T.S., Barch, D.M., Gray, J.R., Molfese, D.L., and Snyder, A. (2001). Anterior Cingulate Cortex and Response Conflict: Effects of Frequency, Inhibition and Errors. Cereb. Cortex 11, 825–836. 10.1093/cercor/11.9.825.

60. Brown, J.W., and Braver, T.S. (2005). Learned Predictions of Error Likelihood in the Anterior Cingulate Cortex. Science 307, 1118–1121. 10.1126/science.1105783.

61. Brockett, A.T., Tennyson, S.S., deBettencourt, C.A., Gaye, F., and Roesch, M.R. (2020). Anterior cingulate cortex is necessary for adaptation of action plans. Proc. Natl. Acad. Sci. 117, 6196–6204. 10.1073/pnas.1919303117.

62. Ito, S., Stuphorn, V., Brown, J.W., and Schall, J.D. (2003). Performance Monitoring by the Anterior Cingulate Cortex During Saccade Countermanding. Science 302, 120–122. 10.1126/science.1087847.

63. Caruana, F., Gerbella, M., Avanzini, P., Gozzo, F., Pelliccia, V., Mai, R., Abdollahi, R.O., Cardinale, F., Sartori, I., Lo Russo, G., et al. (2018). Motor and emotional behaviours elicited by electrical stimulation of the human cingulate cortex. Brain 141, 3035–3051. 10.1093/brain/awy219.

64. Solbakk, A.-K., Funderud, I., Løvstad, M., Endestad, T., Meling, T., Lindgren, M., Knight, R.T., and Krämer, U.M. (2014). Impact of Orbitofrontal Lesions on Electrophysiological Signals in a Stop Signal Task. J. Cogn. Neurosci. 26, 1528–1545. 10.1162/jocn_a_00561.

65. Bryden, D.W., and Roesch, M.R. (2015). Executive Control Signals in Orbitofrontal Cortex during Response Inhibition. J. Neurosci. 35, 3903–3914. 10.1523/JNEUROSCI.3587-14.2015.

66. Ter Horst, J., Boillot, M., Cohen, M.X., and Englitz, B. (2024). Decreased Beta Power and OFC–STN Phase Synchronization during Reactive Stopping in Freely Behaving Rats. J. Neurosci. 44, e0463242024. 10.1523/JNEUROSCI.0463-24.2024.

67. Brockett, A.T., and Roesch, M.R. (2021). The ever-changing OFC landscape: What neural signals in OFC can tell us about inhibitory control. Behav. Neurosci. 135, 129–137. 10.1037/bne0000412.

68. Aponik-Gremillion, L., Chen, Y.Y., Bartoli, E., Koslov, S.R., Rey, H.G., Weiner, K.S., Yoshor, D., Hayden, B.Y., Sheth, S.A., and Foster, B.L. (2022). Distinct population and single-neuron selectivity for executive and episodic processing in human dorsal posterior cingulate. eLife 11, e80722. 10.7554/eLife.80722.

69. Leech, R., and Sharp, D.J. (2014). The role of the posterior cingulate cortex in cognition and disease. Brain 137, 12–32. 10.1093/brain/awt162.

70. Tomiyama, H., Murayama, K., Nemoto, K., Tomita, M., Hasuzawa, S., Mizobe, T., Kato, K., Matsuo, A., Ohno, A., Kan, M., et al. (2023). Posterior cingulate cortex spontaneous activity associated with motor response inhibition in patients with obsessive-compulsive disorder: A resting-state fMRI study. Psychiatry Res. Neuroimaging 334, 111669. 10.1016/j.pscychresns.2023.111669.

71. Foster, B.L., Koslov, S.R., Aponik-Gremillion, L., Monko, M.E., Hayden, B.Y., and Heilbronner, S.R. (2023). A tripartite view of the posterior cingulate cortex. Nat. Rev. Neurosci. 24, 173–189. 10.1038/s41583-022-00661-x.

72. Fox, K.C.R., Foster, B.L., Kucyi, A., Daitch, A.L., and Parvizi, J. (2018). Intracranial Electrophysiology of the Human Default Network. Trends Cogn. Sci. 22, 307–324. 10.1016/j.tics.2018.02.002.

73. Bartoli, E., Aron, A.R., and Tandon, N. (2018). Topography and timing of activity in right inferior frontal cortex and anterior insula for stopping movement. Hum. Brain Mapp. 39, 189–203. 10.1002/hbm.23835.

74. Wessel, J.R., Waller, D.A., and Greenlee, J.D. (2019). Non-selective inhibition of inappropriate motor-tendencies during response-conflict by a fronto-subthalamic mechanism. eLife 8, e42959. 10.7554/eLife.42959.

75. Hannah, R., Muralidharan, V., Sundby, K.K., and Aron, A.R. (2020). Temporally-precise disruption of prefrontal cortex informed by the timing of beta bursts impairs human action-stopping. NeuroImage 222, 117222. 10.1016/j.neuroimage.2020.117222.

76. Muralidharan, V., Aron, A.R., and Schmidt, R. (2022). Transient beta modulates decision thresholds during human action-stopping. NeuroImage 254, 119145. 10.1016/j.neuroimage.2022.119145.

77. Castiglione, A., Wagner, J., Anderson, M., and Aron, A.R. (2019). Preventing a Thought from Coming to Mind Elicits Increased Right Frontal Beta Just as Stopping Action Does. Cereb. Cortex 29, 2160–2172. 10.1093/cercor/bhz017.

78. Errington, S.P., Woodman, G.F., and Schall, J.D. (2020). Dissociation of Medial Frontal β-Bursts and Executive Control. J. Neurosci. 40, 9272–9282. 10.1523/JNEUROSCI.2072-20.2020.

79. Zhang, Y., Chen, Y., Bressler, S.L., and Ding, M. (2008). Response preparation and inhibition: The role of the cortical sensorimotor beta rhythm. Neuroscience 156, 238–246. 10.1016/j.neuroscience.2008.06.061.

80. Schaum, M., Pinzuti, E., Sebastian, A., Lieb, K., Fries, P., Mobascher, A., Jung, P., Wibral, M., and Tüscher, O. (2021). Right inferior frontal gyrus implements motor inhibitory control via beta-band oscillations in humans. eLife 10, e61679. 10.7554/eLife.61679.

81. Verbruggen, F., and Logan, G.D. (2009). Models of response inhibition in the stop-signal and stop-change paradigms. Neurosci. Biobehav. Rev. 33, 647–661. 10.1016/j.neubiorev.2008.08.014.

82. Verbruggen, F., Logan, G.D., and Stevens, M.A. (2008). STOP-IT: Windows executable software for the stop-signal paradigm. Behav. Res. Methods 40, 479–483. 10.3758/brm.40.2.479.

83. Bastos, A.M., and Schoffelen, J.-M. (2016). A Tutorial Review of Functional Connectivity Analysis Methods and Their Interpretational Pitfalls. Front. Syst. Neurosci. 9. 10.3389/fnsys.2015.00175.

84. Schneider, M., Broggini, A.C., Dann, B., Tzanou, A., Uran, C., Sheshadri, S., Scherberger, H., and Vinck, M. (2021). A mechanism for inter-areal coherence through communication based on connectivity and oscillatory power. Neuron 109, 4050–4067.e12. 10.1016/j.neuron.2021.09.037.

85. Strahnen, D., Kapanaiah, S.K.T., Bygrave, A.M., and Kätzel, D. (2021). Lack of redundancy between electrophysiological measures of long-range neuronal communication. BMC Biol. 19, 24. 10.1186/s12915-021-00950-4.

86. Mitzdorf, U. (1985). Current source-density method and application in cat cerebral cortex: investigation of evoked potentials and EEG phenomena. Physiol. Rev. 65, 37–100. 10.1152/physrev.1985.65.1.37.

87. Logothetis, N.K. (2003). The Underpinnings of the BOLD Functional Magnetic Resonance Imaging Signal. J. Neurosci. 23, 3963–3971. 10.1523/JNEUROSCI.23-10-03963.2003.

88. Vinck, M., Battaglia, F.P., Womelsdorf, T., and Pennartz, C. (2012). Improved measures of phase-coupling between spikes and the Local Field Potential. J. Comput. Neurosci. 33, 53–75. 10.1007/s10827-011-0374-4.

89. Unakafova, V.A., and Gail, A. (2019). Comparing Open-Source Toolboxes for Processing and Analysis of Spike and Local Field Potentials Data. Front. Neuroinformatics 13, 57. 10.3389/fninf.2019.00057.

90. Wessel, J.R. (2020). β-Bursts Reveal the Trial-to-Trial Dynamics of Movement Initiation and Cancellation. J. Neurosci. 40, 411–423. 10.1523/JNEUROSCI.1887-19.2019.

91. Haufe, S., Nikulin, V.V., and Nolte, G. (2012). Alleviating the Influence of Weak Data Asymmetries on Granger-Causal Analyses. In Latent Variable Analysis and Signal Separation Lecture Notes in Computer Science., F. Theis, A. Cichocki, A. Yeredor, and M. Zibulevsky, eds. (Springer Berlin Heidelberg), pp. 25–33. 10.1007/978-3-642-28551-6_4.

92. Nogueira, R., Abolafia, J.M., Drugowitsch, J., Balaguer-Ballester, E., Sanchez-Vives, M.V., and Moreno-Bote, R. (2017). Lateral orbitofrontal cortex anticipates choices and integrates prior with current information. Nat. Commun. 8, 14823. 10.1038/ncomms14823.

93. Vinck, M., van Wingerden, M., Womelsdorf, T., Fries, P., and Pennartz, C.M.A. (2010). The pairwise phase consistency: A bias-free measure of rhythmic neuronal synchronization. NeuroImage 51, 112–122. 10.1016/j.neuroimage.2010.01.073.

94. Buckner, R.L., Andrews-Hanna, J.R., and Schacter, D.L. (2008). *The Brain’s Default Network*: *Anatomy, Function, and Relevance to Disease*. Ann. N. Y. Acad. Sci. 1124, 1–38. 10.1196/annals.1440.011.

95. Friston, K.J. (2011). Functional and Effective Connectivity: A Review. Brain Connect. 1, 13–36. 10.1089/brain.2011.0008.

96. Corbetta, M. (2012). Functional connectivity and neurological recovery. Dev. Psychobiol. 54, 239–253. 10.1002/dev.20507.

97. Avena-Koenigsberger, A., Misic, B., and Sporns, O. (2018). Communication dynamics in complex brain networks. Nat. Rev. Neurosci. 19, 17–33. 10.1038/nrn.2017.149.

98. Menon, V. (2011). Large-scale brain networks and psychopathology: a unifying triple network model. Trends Cogn. Sci. 15, 483–506. 10.1016/j.tics.2011.08.003.

99. Baker, J.T., Holmes, A.J., Masters, G.A., Yeo, B.T.T., Krienen, F., Buckner, R.L., and Öngür, D. (2014). Disruption of Cortical Association Networks in Schizophrenia and Psychotic Bipolar Disorder. JAMA Psychiatry 71, 109. 10.1001/jamapsychiatry.2013.3469.

100. Fornito, A., Zalesky, A., and Breakspear, M. (2015). The connectomics of brain disorders. Nat. Rev. Neurosci. 16, 159–172. 10.1038/nrn3901.

101. Lynch, C.J., Elbau, I.G., Ng, T., Ayaz, A., Zhu, S., Wolk, D., Manfredi, N., Johnson, M., Chang, M., Chou, J., et al. (2024). Frontostriatal salience network expansion in individuals in depression. Nature. 10.1038/s41586-024-07805-2.

102. Siegel, J.S., Subramanian, S., Perry, D., Kay, B.P., Gordon, E.M., Laumann, T.O., Reneau, T.R., Metcalf, N.V., Chacko, R.V., Gratton, C., et al. (2024). Psilocybin desynchronizes the human brain. Nature 632, 131–138. 10.1038/s41586-024-07624-5.

103. Wang, B.A., Veismann, M., Banerjee, A., and Pleger, B. (2023). Human orbitofrontal cortex signals decision outcomes to sensory cortex during behavioral adaptations. Nat. Commun. 14, 3552. 10.1038/s41467-023-38671-7.

104. Morecraft, R.J., Stilwell-Morecraft, K.S., Cipolloni, P.B., Ge, J., McNeal, D.W., and Pandya, D.N. (2012). Cytoarchitecture and cortical connections of the anterior cingulate and adjacent somatomotor fields in the rhesus monkey. Brain Res. Bull. 87, 457–497. 10.1016/j.brainresbull.2011.12.005.

105. Cavada, C., Compañy, T., Tejedor, J., Cruz-Rizzolo, R.J., and Reinoso-Suárez, F. (2000). The Anatomical Connections of the Macaque Monkey Orbitofrontal Cortex. A Review. Cereb. Cortex 10, 220–242. 10.1093/cercor/10.3.220.

106. Jahanshahi, M., Obeso, I., Rothwell, J.C., and Obeso, J.A. (2015). A fronto–striato– subthalamic–pallidal network for goal-directed and habitual inhibition. Nat. Rev. Neurosci. 16, 719–732. 10.1038/nrn4038.

107. Diesburg, D.A., Greenlee, J.D., and Wessel, J.R. (2021). Cortico-subcortical β burst dynamics underlying movement cancellation in humans. eLife 10, e70270. 10.7554/eLife.70270.

108. Andersen, R.A., and Cui, H. (2009). Intention, Action Planning, and Decision Making in Parietal-Frontal Circuits. Neuron 63, 568–583. 10.1016/j.neuron.2009.08.028.

109. Power, J.D., Cohen, A.L., Nelson, S.M., Wig, G.S., Barnes, K.A., Church, J.A., Vogel, A.C., Laumann, T.O., Miezin, F.M., Schlaggar, B.L., et al. (2011). Functional Network Organization of the Human Brain. Neuron 72, 665–678. 10.1016/j.neuron.2011.09.006.

110. Selemon, L., and Goldman-Rakic, P. (1988). Common cortical and subcortical targets of the dorsolateral prefrontal and posterior parietal cortices in the rhesus monkey: evidence for a distributed neural network subserving spatially guided behavior. J. Neurosci. 8, 4049–4068. 10.1523/JNEUROSCI.08-11-04049.1988.

111. Gerbella, M., Borra, E., Tonelli, S., Rozzi, S., and Luppino, G. (2013). Connectional Heterogeneity of the Ventral Part of the Macaque Area 46. Cereb. Cortex 23, 967–987. 10.1093/cercor/bhs096.

112. Logothetis, N.K. (2008). What we can do and what we cannot do with fMRI. Nature 453, 869–878. 10.1038/nature06976.

113. Romero, M.C., Davare, M., Armendariz, M., and Janssen, P. (2019). Neural effects of transcranial magnetic stimulation at the single-cell level. Nat. Commun. 10, 2642. 10.1038/s41467-019-10638-7.

114. Siebner, H.R., Funke, K., Aberra, A.S., Antal, A., Bestmann, S., Chen, R., Classen, J., Davare, M., Di Lazzaro, V., Fox, P.T., et al. (2022). Transcranial magnetic stimulation of the brain: What is stimulated? – A consensus and critical position paper. Clin. Neurophysiol. 140, 59–97. 10.1016/j.clinph.2022.04.022.

115. Guan, Y., and Wessel, J.R. (2022). Two Types of Motor Inhibition after Action Errors in Humans. J. Neurosci. 42, 7267–7275. 10.1523/JNEUROSCI.1191-22.2022.

116. Balasubramani, P.P., Pesce, M.C., and Hayden, B.Y. (2020). Activity in orbitofrontal neuronal ensembles reflects inhibitory control. Eur. J. Neurosci. 51, 2033–2051. 10.1111/ejn.14638.

117. Yoo, S.B.M., and Hayden, B.Y. (2018). Economic Choice as an Untangling of Options into Actions. Neuron 99, 434–447. 10.1016/j.neuron.2018.06.038.

118. Wallis, J.D. (2007). Orbitofrontal Cortex and Its Contribution to Decision-Making. Annu. Rev. Neurosci. 30, 31–56. 10.1146/annurev.neuro.30.051606.094334.

119. Padoa-Schioppa, C., and Conen, K.E. (2017). Orbitofrontal Cortex: A Neural Circuit for Economic Decisions. Neuron 96, 736–754. 10.1016/j.neuron.2017.09.031.

120. Schoenbaum, G., and Roesch, M. (2005). Orbitofrontal Cortex, Associative Learning, and Expectancies. Neuron 47, 633–636. 10.1016/j.neuron.2005.07.018.

121. Kerns, J.G., Cohen, J.D., MacDonald, A.W., Cho, R.Y., Stenger, V.A., and Carter, C.S. (2004). Anterior Cingulate Conflict Monitoring and Adjustments in Control. Science 303, 1023–1026. 10.1126/science.1089910.

122. Botvinick, M.M., Cohen, J.D., and Carter, C.S. (2004). Conflict monitoring and anterior cingulate cortex: an update. Trends Cogn. Sci. 8, 539–546. 10.1016/j.tics.2004.10.003.

123. Sheth, S.A., Mian, M.K., Patel, S.R., Asaad, W.F., Williams, Z.M., Dougherty, D.D., Bush, G., and Eskandar, E.N. (2012). Human dorsal anterior cingulate cortex neurons mediate ongoing behavioural adaptation. Nature 488, 218–221. 10.1038/nature11239.

124. Bartoli, E., Conner, C.R., Kadipasaoglu, C.M., Yellapantula, S., Rollo, M.J., Carter, C.S., and Tandon, N. (2018). Temporal Dynamics of Human Frontal and Cingulate Neural Activity During Conflict and Cognitive Control. Cereb. Cortex 28, 3842–3856. 10.1093/cercor/bhx245.

125. Isherwood, S.J.S., Keuken, M.C., Bazin, P.L., and Forstmann, B.U. (2021). Cortical and subcortical contributions to interference resolution and inhibition – An fMRI ALE meta-analysis. Neurosci. Biobehav. Rev. 129, 245–260. 10.1016/j.neubiorev.2021.07.021.

126. Heilbronner, S.R., and Hayden, B.Y. (2016). Dorsal Anterior Cingulate Cortex: A Bottom-Up View. Annu. Rev. Neurosci. 39, 149–170. 10.1146/annurev-neuro-070815-013952.

127. Hayden, B.Y., and Platt, M.L. (2010). Neurons in Anterior Cingulate Cortex Multiplex Information about Reward and Action. J. Neurosci. 30, 3339–3346. 10.1523/JNEUROSCI.4874-09.2010.

128. Pourtois, G., Vocat, R., N’Diaye, K., Spinelli, L., Seeck, M., and Vuilleumier, P. (2010). Errors recruit both cognitive and emotional monitoring systems: Simultaneous intracranial recordings in the dorsal anterior cingulate gyrus and amygdala combined with fMRI. Neuropsychologia 48, 1144–1159. 10.1016/j.neuropsychologia.2009.12.020.

129. Dosenbach, N.U.F., Raichle, M.E., and Gordon, E.M. (2025). The brain’s action-mode network. Nat. Rev. Neurosci., 1–11. 10.1038/s41583-024-00895-x.

130. Raichle, M.E. (2010). Two views of brain function. Trends Cogn. Sci. 14, 180–190. 10.1016/j.tics.2010.01.008.

131. Li, C.-S.R., Yan, P., Bergquist, K.L., and Sinha, R. (2007). Greater activation of the “default” brain regions predicts stop signal errors. NeuroImage 38, 640–648. 10.1016/j.neuroimage.2007.07.021.

132. Heilbronner, S.R., and Platt, M.L. (2013). Causal Evidence of Performance Monitoring by Neurons in Posterior Cingulate Cortex during Learning. Neuron 80, 1384–1391. 10.1016/j.neuron.2013.09.028.

133. Hayden, B.Y., Nair, A.C., McCoy, A.N., and Platt, M.L. (2008). Posterior Cingulate Cortex Mediates Outcome-Contingent Allocation of Behavior. Neuron 60, 19–25. 10.1016/j.neuron.2008.09.012.

134. Hellyer, P.J., Shanahan, M., Scott, G., Wise, R.J.S., Sharp, D.J., and Leech, R. (2014). The Control of Global Brain Dynamics: Opposing Actions of Frontoparietal Control and Default Mode Networks on Attention. J. Neurosci. 34, 451–461. 10.1523/JNEUROSCI.1853-13.2014.

135. Menon, V. (2023). 20 years of the default mode network: A review and synthesis. Neuron 111, 2469–2487. 10.1016/j.neuron.2023.04.023.

136. Shofty, B., Gonen, T., Bergmann, E., Mayseless, N., Korn, A., Shamay-Tsoory, S., Grossman, R., Jalon, I., Kahn, I., and Ram, Z. (2022). The default network is causally linked to creative thinking. Mol. Psychiatry 27, 1848–1854. 10.1038/s41380-021-01403-8.

137. Bartoli, E., Devara, E., Dang, H.Q., Rabinovich, R., Mathura, R.K., Anand, A., Pascuzzi, B.R., Adkinson, J., Kenett, Y.N., Bijanki, K.R., et al. (2024). Default mode network electrophysiological dynamics and causal role in creative thinking. Brain 147, 3409–3425. 10.1093/brain/awae199.

138. Weissman, D.H., Roberts, K.C., Visscher, K.M., and Woldorff, M.G. (2006). The neural bases of momentary lapses in attention. Nat. Neurosci. 9, 971–978. 10.1038/nn1727.

139. Leech, R., and Smallwood, J. (2019). The posterior cingulate cortex: Insights from structure and function. In Handbook of Clinical Neurology (Elsevier), pp. 73–85. 10.1016/B978-0-444-64196-0.00005-4.

140. Heilbronner, S.R., Hayden, B.Y., and Platt, M.L. (2011). Decision Salience Signals in Posterior Cingulate Cortex. Front. Neurosci. 5. 10.3389/fnins.2011.00055.

141. Corlett, P.R., Mollick, J.A., and Kober, H. (2022). Meta-analysis of human prediction error for incentives, perception, cognition, and action. Neuropsychopharmacology 47, 1339– 1349. 10.1038/s41386-021-01264-3.

142. Bolognini, N., and Miniussi, C. (2018). Noninvasive brain stimulation of the parietal lobe for improving neurologic, neuropsychologic, and neuropsychiatric deficits. In Handbook of Clinical Neurology (Elsevier), pp. 427–446. 10.1016/B978-0-444-63622-5.00022-X.

143. MacDonald, P.A., and Paus, T. (2003). The Role of Parietal Cortex in Awareness of Self-generated Movements: a Transcranial Magnetic Stimulation Study. Cereb. Cortex 13, 962– 967. 10.1093/cercor/13.9.962.

144. Iacoboni, M. (2006). Visuo-motor integration and control in the human posterior parietal cortex: Evidence from TMS and fMRI. Neuropsychologia 44, 2691–2699. 10.1016/j.neuropsychologia.2006.04.029.

145. Mevorach, C., Humphreys, G.W., and Shalev, L. (2006). Opposite biases in salience-based selection for the left and right posterior parietal cortex. Nat. Neurosci. 9, 740–742. 10.1038/nn1709.

146. Romei, V., Driver, J., Schyns, P.G., and Thut, G. (2011). Rhythmic TMS over Parietal Cortex Links Distinct Brain Frequencies to Global versus Local Visual Processing. Curr. Biol. 21, 334–337. 10.1016/j.cub.2011.01.035.

147. Lee, J., Ku, J., Han, K., Park, J., Lee, H., Kim, K.R., Lee, E., Husain, M., Yoon, K.J., Kim, I.Y., et al. (2013). rTMS over bilateral inferior parietal cortex induces decrement of spatial sustained attention. Front. Hum. Neurosci. 7. 10.3389/fnhum.2013.00026.

148. Rushworth, M.F.S., and Taylor, P.C.J. (2006). TMS in the parietal cortex: Updating representations for attention and action. Neuropsychologia 44, 2700–2716. 10.1016/j.neuropsychologia.2005.12.007.

149. Jia, Y., Xu, L., Yang, K., Zhang, Y., Lv, X., Zhu, Z., Chen, Z., Zhu, Y., Wei, L., Li, X., et al. (2021). Precision Repetitive Transcranial Magnetic Stimulation Over the Left Parietal Cortex Improves Memory in Alzheimer’s Disease: A Randomized, Double-Blind, Sham-Controlled Study. Front. Aging Neurosci. 13, 693611. 10.3389/fnagi.2021.693611.

150. Prime, S.L., Vesia, M., and Crawford, J.D. (2008). Transcranial Magnetic Stimulation over Posterior Parietal Cortex Disrupts Transsaccadic Memory of Multiple Objects. J. Neurosci. 28, 6938–6949. 10.1523/JNEUROSCI.0542-08.2008.

151. Guo, X., Zhou, Q., Lu, Y., Xu, Z., Wen, Z., Gu, P., Tian, S., and Wang, Y. (2024). Transcranial direct current stimulation over the right parietal cortex improves the depressive disorder: A preliminary study. Brain Behav. 14, e3638. 10.1002/brb3.3638.

152. Huang, Z., Li, Y., Bianchi, M.T., Zhan, S., Jiang, F., Li, N., Ding, Y., Hou, Y., Wang, L., Ouyang, Q., et al. (2018). Repetitive transcranial magnetic stimulation of the right parietal cortex for comorbid generalized anxiety disorder and insomnia: A randomized, double-blind, sham-controlled pilot study. Brain Stimulat. 11, 1103–1109. 10.1016/j.brs.2018.05.016.

153. Balderston, N.L., Beydler, E.M., Goodwin, M., Deng, Z.-D., Radman, T., Luber, B., Lisanby, S.H., Ernst, M., and Grillon, C. (2020). Low-frequency parietal repetitive transcranial magnetic stimulation reduces fear and anxiety. Transl. Psychiatry 10, 68. 10.1038/s41398-020-0751-8.

154. Balderston, N.L., Hale, E., Hsiung, A., Torrisi, S., Holroyd, T., Carver, F.W., Coppola, R., Ernst, M., and Grillon, C. (2017). Threat of shock increases excitability and connectivity of the intraparietal sulcus. eLife 6, e23608. 10.7554/eLife.23608.

155. Siddiqi, S.H., Kording, K.P., Parvizi, J., and Fox, M.D. (2022). Causal mapping of human brain function. Nat. Rev. Neurosci. 23, 361–375. 10.1038/s41583-022-00583-8.

156. Brainard, D.H. (1997). The Psychophysics Toolbox. Spat. Vis. 10, 433–436.

157. Groppe, D.M., Bickel, S., Dykstra, A.R., Wang, X., Mégevand, P., Mercier, M.R., Lado, F.A., Mehta, A.D., and Honey, C.J. (2017). iELVis: An open source MATLAB toolbox for localizing and visualizing human intracranial electrode data. J. Neurosci. Methods 281, 40–48. 10.1016/j.jneumeth.2017.01.022.

158. Jenkinson, M., Beckmann, C.F., Behrens, T.E.J., Woolrich, M.W., and Smith, S.M. (2012). FSL. NeuroImage 62, 782–790. 10.1016/j.neuroimage.2011.09.015.

159. Papademetris, X., Jackowski, M., Rajeevan, N., DiStasio, M., Okuda, H., Constable, R.T., and Staib, L. (2006). BioImage Suite: An integrated medical image analysis suite: An update. Insight J. 10.54294/2g80r4.

160. Dale, A.M., Fischl, B., and Sereno, M.I. (1999). Cortical Surface-Based Analysis: I. Segmentation and Surface Reconstruction. NeuroImage 9, 179–194. 10.1006/nimg.1998.0395.

161. Destrieux, C., Fischl, B., Dale, A., and Halgren, E. (2010). Automatic parcellation of human cortical gyri and sulci using standard anatomical nomenclature. NeuroImage 53, 1–15. 10.1016/j.neuroimage.2010.06.010.

162. Stone, S., Madsen, J.R., Bolton, J., Pearl, P.L., Chavakula, V., and Day, E. (2021). A Standardized Electrode Nomenclature for Stereoelectroencephalography Applications. J. Clin. Neurophysiol. 38, 509–515. 10.1097/WNP.0000000000000724.

163. Oostenveld, R., Fries, P., Maris, E., and Schoffelen, J.-M. (2011). FieldTrip: Open Source Software for Advanced Analysis of MEG, EEG, and Invasive Electrophysiological Data. Comput. Intell. Neurosci. 2011, 1–9. 10.1155/2011/156869.

164. Seth, A.K., Barrett, A.B., and Barnett, L. (2015). Granger Causality Analysis in Neuroscience and Neuroimaging. J. Neurosci. 35, 3293–3297. 10.1523/JNEUROSCI.4399-14.2015.

165. Dhamala, M., Rangarajan, G., and Ding, M. (2008). Analyzing information flow in brain networks with nonparametric Granger causality. NeuroImage 41, 354–362. 10.1016/j.neuroimage.2008.02.020.

166. Mitra, P.P., and Pesaran, B. (1999). Analysis of Dynamic Brain Imaging Data. Biophys. J. 76, 691–708. 10.1016/S0006-3495(99)77236-X.

167. Geweke, J. (1982). Measurement of Linear Dependence and Feedback between Multiple Time Series. J. Am. Stat. Assoc. 77, 304–313. 10.1080/01621459.1982.10477803.

168. Srinath, R., and Ray, S. (2014). Effect of amplitude correlations on coherence in the local field potential. J. Neurophysiol. 112, 741–751. 10.1152/jn.00851.2013.

169. Chaure, F.J., Rey, H.G., and Quian Quiroga, R. (2018). A novel and fully automatic spike-sorting implementation with variable number of features. J. Neurophysiol. 120, 1859– 1871. 10.1152/jn.00339.2018.

170. Rey, H.G., Gori, B., Chaure, F.J., Collavini, S., Blenkmann, A.O., Seoane, P., Seoane, E., Kochen, S., and Quian Quiroga, R. (2020). Single Neuron Coding of Identity in the Human Hippocampal Formation. Curr. Biol. 30, 1152–1159.e3. 10.1016/j.cub.2020.01.03

