## Supplemental Information for "Parietal cortex is recruited by frontal and cingulate areas to support action monitoring and updating during stopping"

| **Participant information** | | | | | | | | | |
|---|---|---|---|---|---|---|---|---|---|
| **#** | **Age** | **Sex** | **Posterior parietal cortex** | | **SSRT** | **p(resp\|stop)** | **Inclusion in analysis** | | |
| | | | **R** | **L** | | | **TF** | **LFP-LFP** | **spike-LFP** |
| **1** | 53 | F | 13 | - | 330 | 0.50 | y | y | - |
| **2** | 37 | M | 1 | 5 | 134 | 0.41 | y | - | - |
| **3** | 44 | F | 4 | 3 | 218 | 0.49 | y | - | - |
| **4** | 39 | F | - | 9 | 211 | 0.46 | y | y | - |
| **5** | 25 | M | - | 8 | 252 | 0.47 | y | y | - |
| **6** | 30 | M | - | 4 | 199 | 0.39 | y | - | y |
| **7** | 24 | M | 8 | 1 | 120 | 0.33 | y | y | y |
| **8** | 32 | F | 15 | 5 | 316 | 0.51 | y | y | y |
| **9** | 33 | F | 13 | 24 | 324 | 0.49 | y | y | y |
| **10** | 52 | M | 11 | 17 | 252 | 0.48 | y | y | y |
| **11** | 25 | F | - | 2 | 245 | 0.39 | y | - | y |
| **12** | 39 | M | 9 | - | 325 | 0.51 | y | - | - |

Table S1. Patient demographics, number of intracranial electrodes in posterior parietal cortex, behavioral data (SSRT and accuracy), and details for inclusion in time-frequency (TF), LFP-LFP, and spike-LFP analyses

| | LFP-LFP Granger causality and PPC | | | | | | | Spike-LFP PPC | | |
|---|---|---|---|---|---|---|---|---|---|---|
| **#** | **IPS** | **IFG** | **ACC** | **PCC** | **OFC** | **R-IPS** | **R-IFG** | **ACC (SUA)** | **PCC (SUA)** | **Parietal (LFP)** |
| **1** | 4 | 4 | 3 | - | 25 | 4 | 4 | - | - | - |
| **4** | 3 | 5 | 4 | 2 | 6 | - | - | - | - | - |
| **5** | 2 | 1 | 24 | 10 | 4 | - | - | - | - | - |
| **6** | - | - | - | - | - | - | - | 2 | 4 | 4 |
| **7** | 2 | 13 | 16 | - | 20 | 2 | 5 | 7 | - | 9 |
| **8** | 10 | 3 | 2 | 2 | 3 | 9 | 1 | 2 | 2 | 20 |
| **9** | 17 | 1 | 2 | 11 | - | 11 | 1 | 28 | - | 37 |
| **10** | 3 | - | 6 | - | - | - | - | 13 | - | 38 |
| **11** | - | - | - | - | - | - | - | 10 | 6 | 2 |
| ***Sum*** | ***41*** | ***27*** | ***57*** | ***25*** | ***58*** | ***26*** | ***11*** | ***62*** | ***12*** | ***110*** |

Table S2: Intracranial electrodes in each patient that were used in functional connectivity analysis

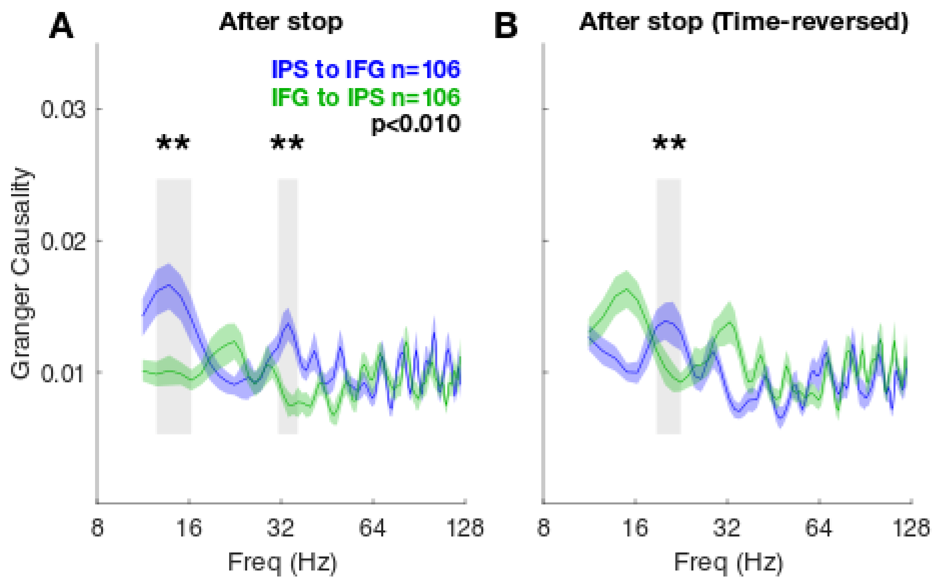


Figure S1. Time-reversal analysis of LFP-LFP Granger causality between IPS and IFG

Time-reversal of original data should invert the order of Granger causality between two signals. If the order does not invert, this is likely due to external noise or common input.[83,91] In both panels, the black asterisks and gray vertical shaded regions denote $p < 0.01$ (Wilcoxon signed-rank test), and colored shaded regions denote ± SEM.

(A) Granger causality from original data in the epoch [0 800 ms] aligned to stop cue onset. Granger causality from IPS to IFG is significantly higher than vice versa at 12-16 and 31-36 Hz.

(B) Granger causality from time-reversed data in the epoch [0 800 ms] aligned to stop cue onset. Granger causality from IPS to IFG is significantly higher than vice versa at 19-23 Hz. Peak IFG-to-IPS Granger causality was 0.016 at 15 Hz, while peak IPS-to-IFG Granger causality was 0.014 at 19 Hz. This suggests that the observed effect in (A) is reliable rather than due to external noise. The inversion effect after time reversal was not significant in frequencies below 10 Hz (data not shown).

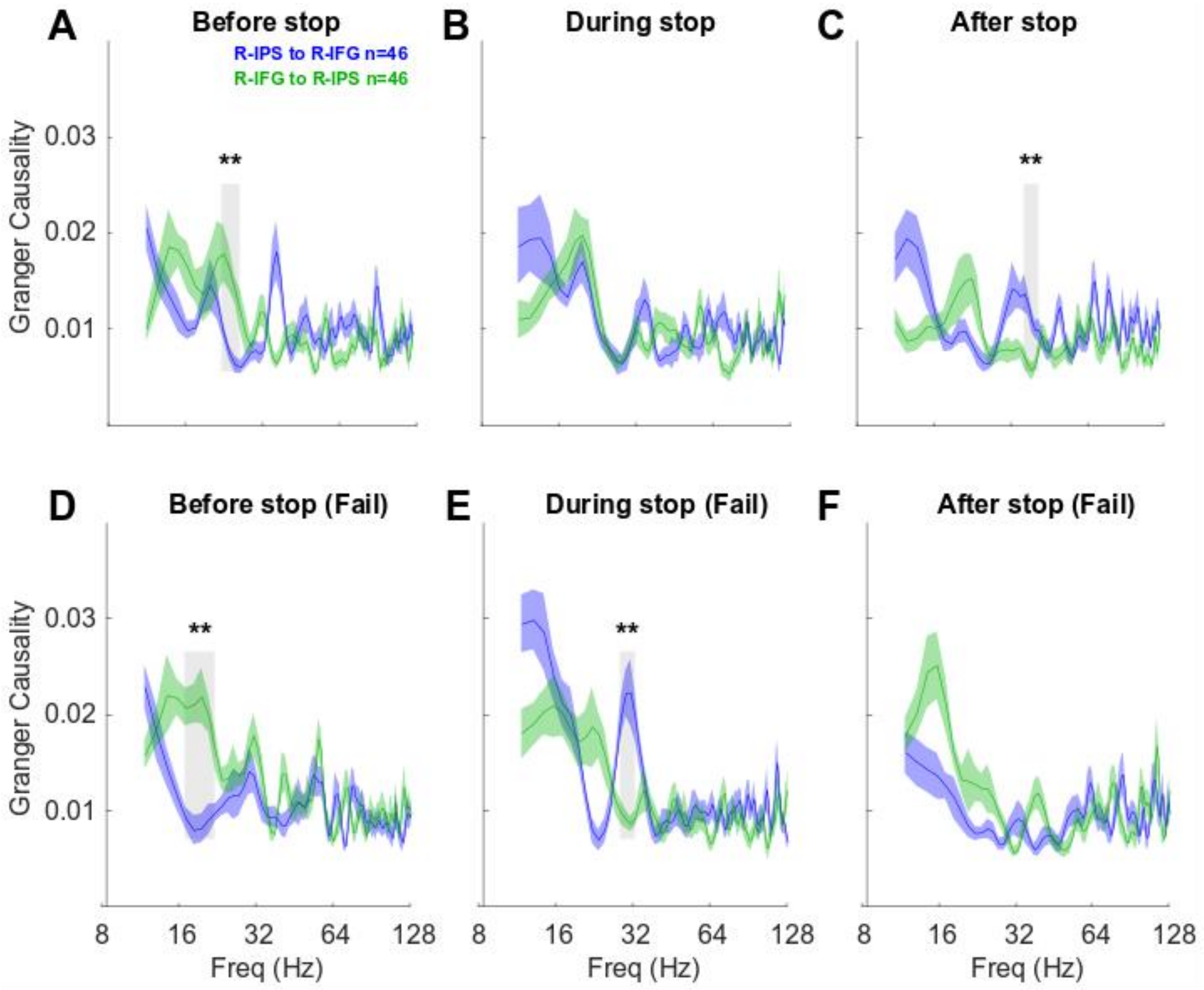


Figure S2. Granger causality between right IPS and right IFG in Stop-success and Stop-fail trials

In all panels, black asterisks and gray vertical shaded regions denote $p < 0.01$ (Wilcoxon signed-rank test).

(A-C) Granger causality in Stop-success trials. Format same as Figure 2A-C. (D-F) Granger causality in Stop-fail trials. Format same as Figure 2D-F.

(A) In the Before-stop epoch, Granger causality is significantly higher from IFG to IPS than vice versa at 23-26 Hz.

(B) In the During-stop epoch, there is no significant difference between Granger causality from IFG to IPS and vice versa.

(C) In the After-stop epoch, Granger causality is significantly higher from IPS to IFG than vice versa at 36-41 Hz.

(D) In the Before-stop epoch, Granger causality is significantly higher from IFG to IPS than vice versa at 16-21 Hz.

(E) In the During-stop epoch, Granger causality is significantly higher from IPS to IFG than vice versa at 28-31 Hz.

(F) In the After-stop epoch, there is no significant difference between Granger causality from IFG to IPS and vice versa.

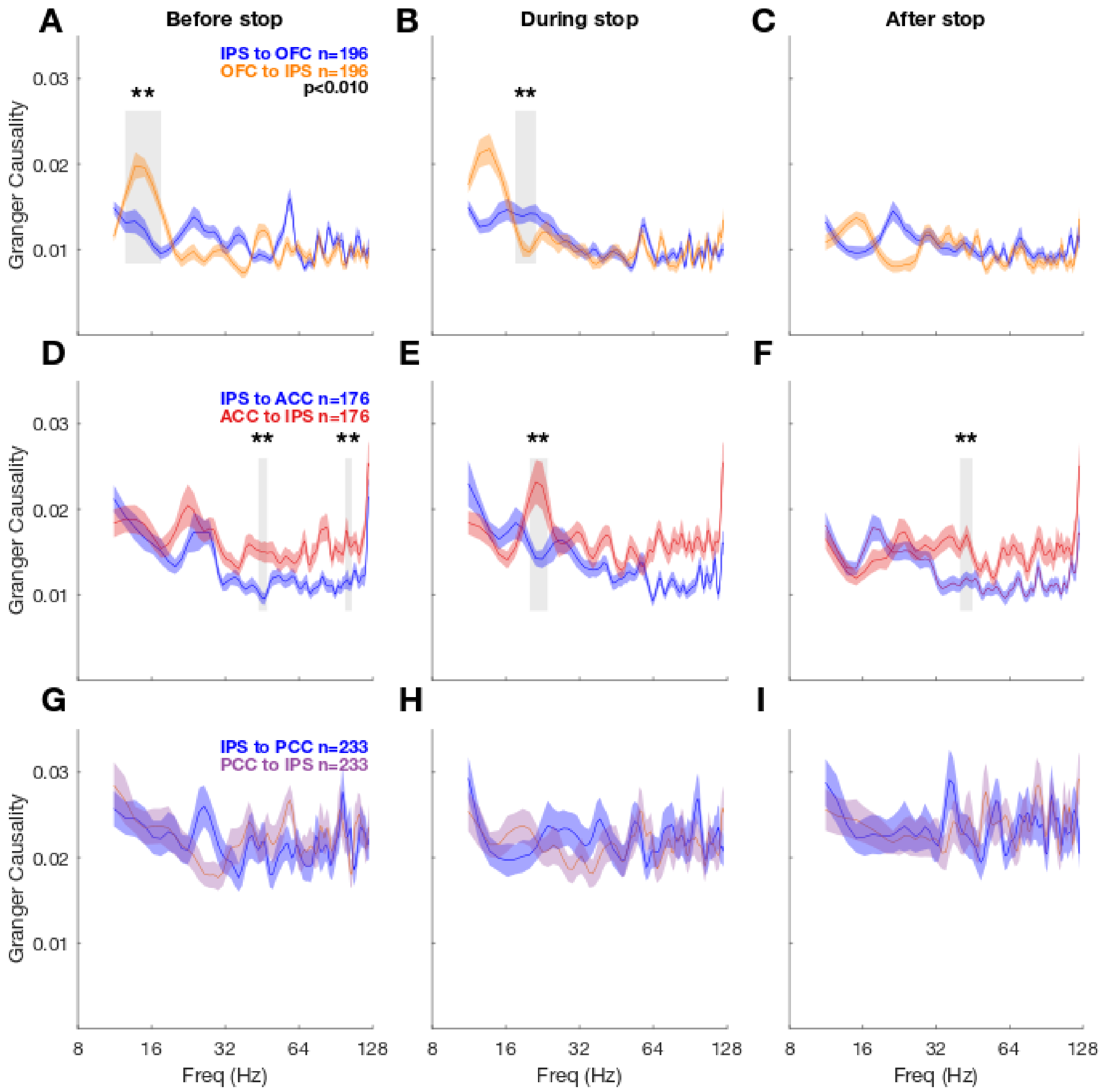


### Figure S3. Granger causality in IPS-OFC, IPS-ACC, and IPS-PCC across different time periods in successful movement inhibition (Stop-success)

Same format as in Figure 2A-C. Black asterisks and gray vertical shaded regions denote $p < 0.01$ (Wilcoxon signed-rank test).

(A), (B), and (C): Granger causality between IPS and OFC. There is asymmetry in Granger causality values in beta frequency range.

(D), (E), and (F): Granger causality between IPS and ACC. There is asymmetry in Granger causality values in beta and gamma frequency range.

(G), (H), and (I): Granger causality between IPS and PCC. There is no significant difference between Granger causality from PCC to IPS and vice versa.

(A) In the Before-stop epoch, Granger causality is significantly higher from OFC to IPS than vice versa at 13-18 Hz.

(B) In the During-stop epoch, Granger causality is significantly higher from IPS to OFC than vice versa at 18-21 Hz.

(C) In the After-stop epochs, there is no significant difference between Granger causality from OFC to IPS and vice versa.

(D) In the Before-stop epoch, Granger causality is significantly higher from ACC to IPS than vice versa at 44-48 and 99-105 Hz.

(E) In the During-stop epoch, Granger causality is significantly higher from ACC to IPS than vice versa at 20-24 Hz.

(F) In the After-stop epoch, Granger causality is significantly higher from ACC to IPS than vice versa at 40-45 Hz.

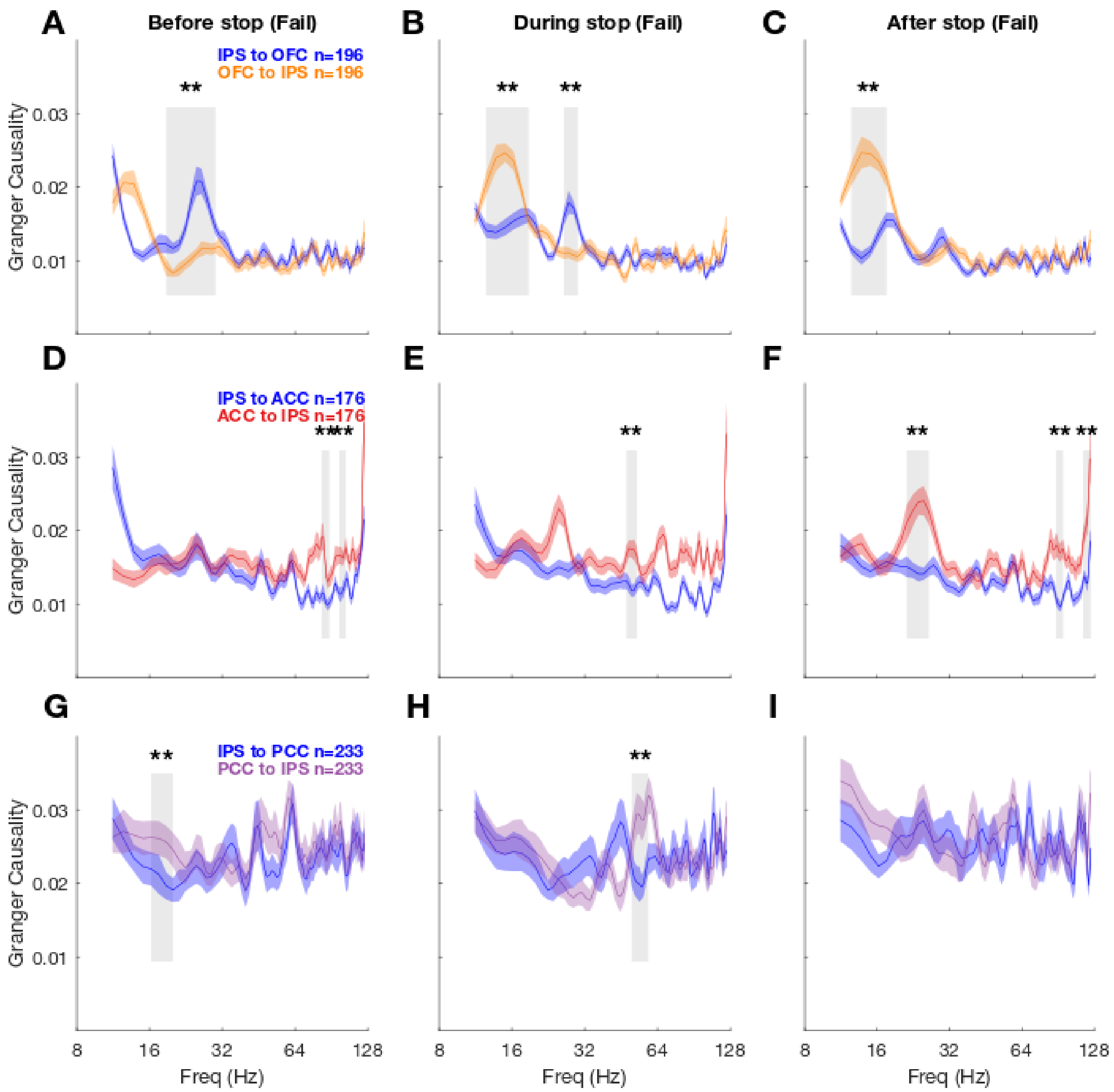


Figure S4. Granger causality in IPS-OFC, IPS-ACC, IPS-PCC across different time periods in stop error trials (Stop-fail)

Same format as in Figure 2D-F. Black asterisks and gray vertical shaded regions denote $p < 0.01$ (Wilcoxon signed-rank test).

(A), (B), and (C): Granger causality between IPS and OFC.

(D), (E), and (F): Granger causality between IPS and ACC. There is asymmetry in Granger causality values in beta and high gamma frequency range.

(G), (H), and (I): Granger causality between IPS and PCC.

(A) In the Before-stop epoch, Granger causality from IPS to OFC is significantly higher than vice versa at 19-30 Hz.

(B) In the During-stop epoch, Granger causality is significantly higher from OFC to IPS than vice versa at 13-19 Hz. However, Granger causality is significantly higher from IPS to OFC than vice versa at 26-30 Hz.

(C) In the After-stop epochs, Granger causality is significantly higher from OFC to IPS than vice versa at 13-18 Hz.

(D) In the Before-stop epoch, Granger causality is significantly higher from ACC to IPS than vice versa at 83-89 Hz and 98-104 Hz.

(E) In the During-stop epoch, Granger causality is significantly higher from ACC to IPS than vice versa at 48-53 Hz.

(F) In the After-stop epoch, Granger causality is significantly higher from ACC to IPS than vice versa at 21-26, 89-95, and 115-124 Hz.

(G) In the Before-stop epoch, Granger causality is significantly higher from PCC to IPS than vice versa at 16-20 Hz.

(H) In the During-stop epoch, Granger causality is significantly higher from PCC to IPS than vice versa at 50-59 Hz.

(I) In the After-stop epoch, there is no significant difference between Granger causality from PCC to IPS and vice versa.

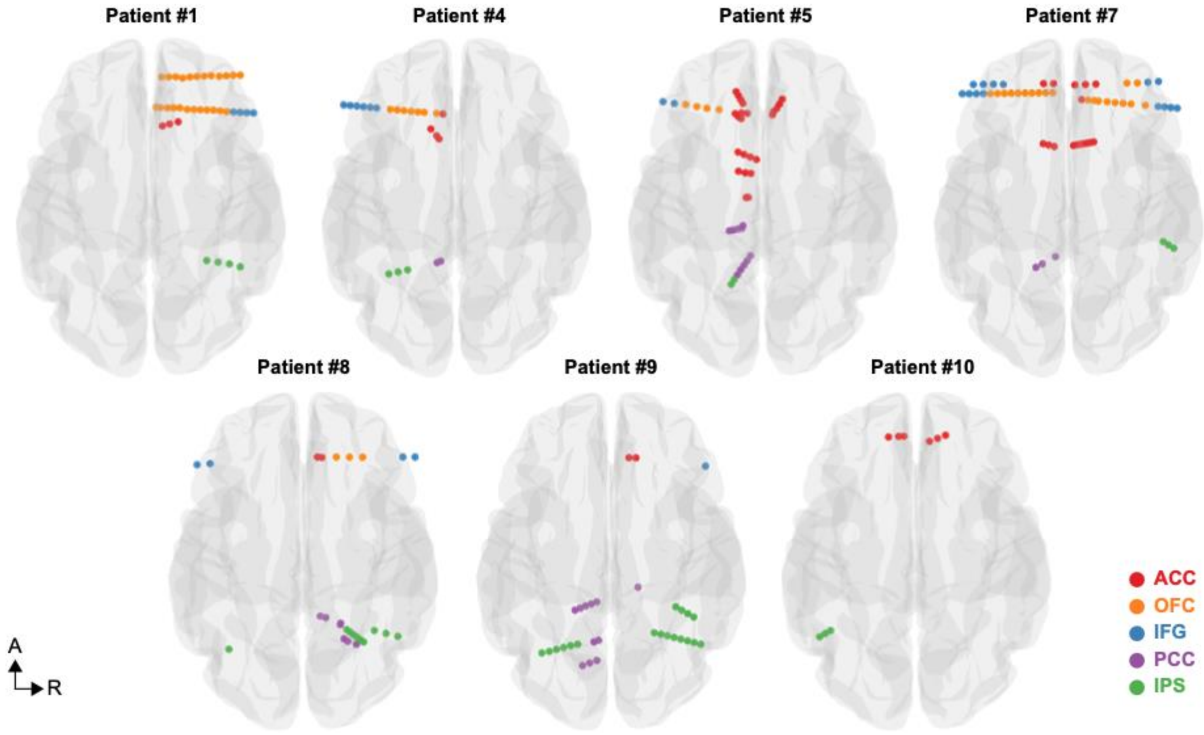


### Figure S5. Intracranial recording locations from individual subjects used for connectivity analyses

Intracranial recording locations in ACC, IFG, IPS, OFC, and PCC shown separately for each of the 7 subjects for the Granger causality and LFP-LFP PPC analysis. Number of intracranial recording contacts are shown in Table S2.

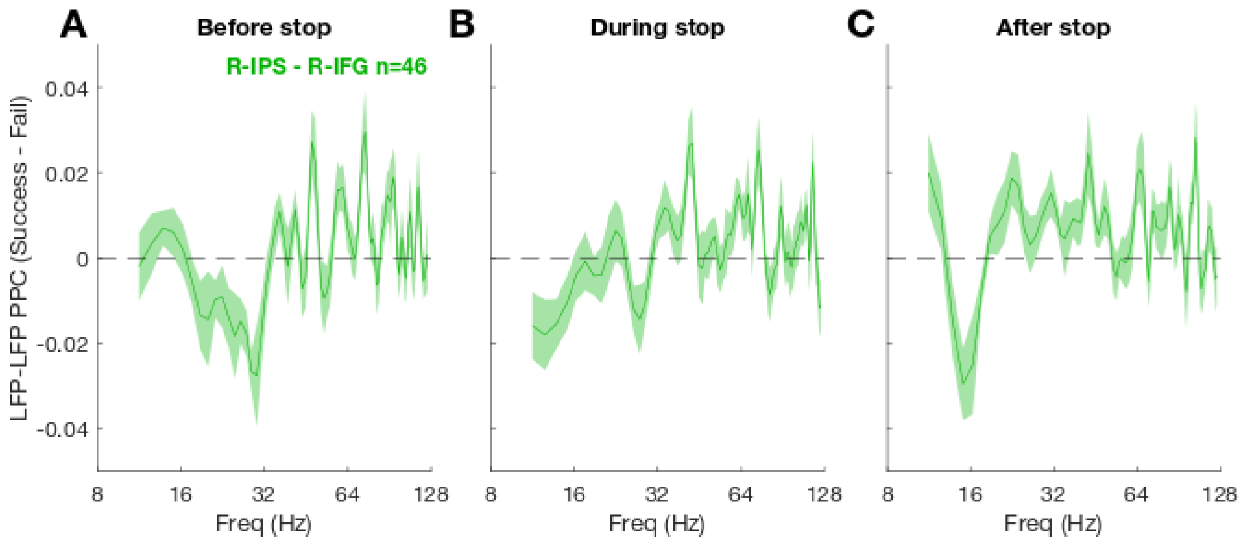


### Figure S6. LFP-LFP PPC between right IPS and right IFG

Same format as Figure 5A-C. We measured LFP-LFP PPC between right IPS and right IFG (4 patients). There is no significant modulation in 31-35 Hz in the During-stop epoch in (B) that was previously observed in PPC between all IPS and IFG LFP pairs in Figure 5B. However, the PPC difference between success and fail trials is positive in (B) similar to the results from all IPS and IFG LFP pairs. The difference in (B) is statistically significant at individual frequency data points at 31, 34, and 35 Hz ($p < 0.01$, paired t-test), thus failing to reach our criterion of four contiguous significant frequency values.

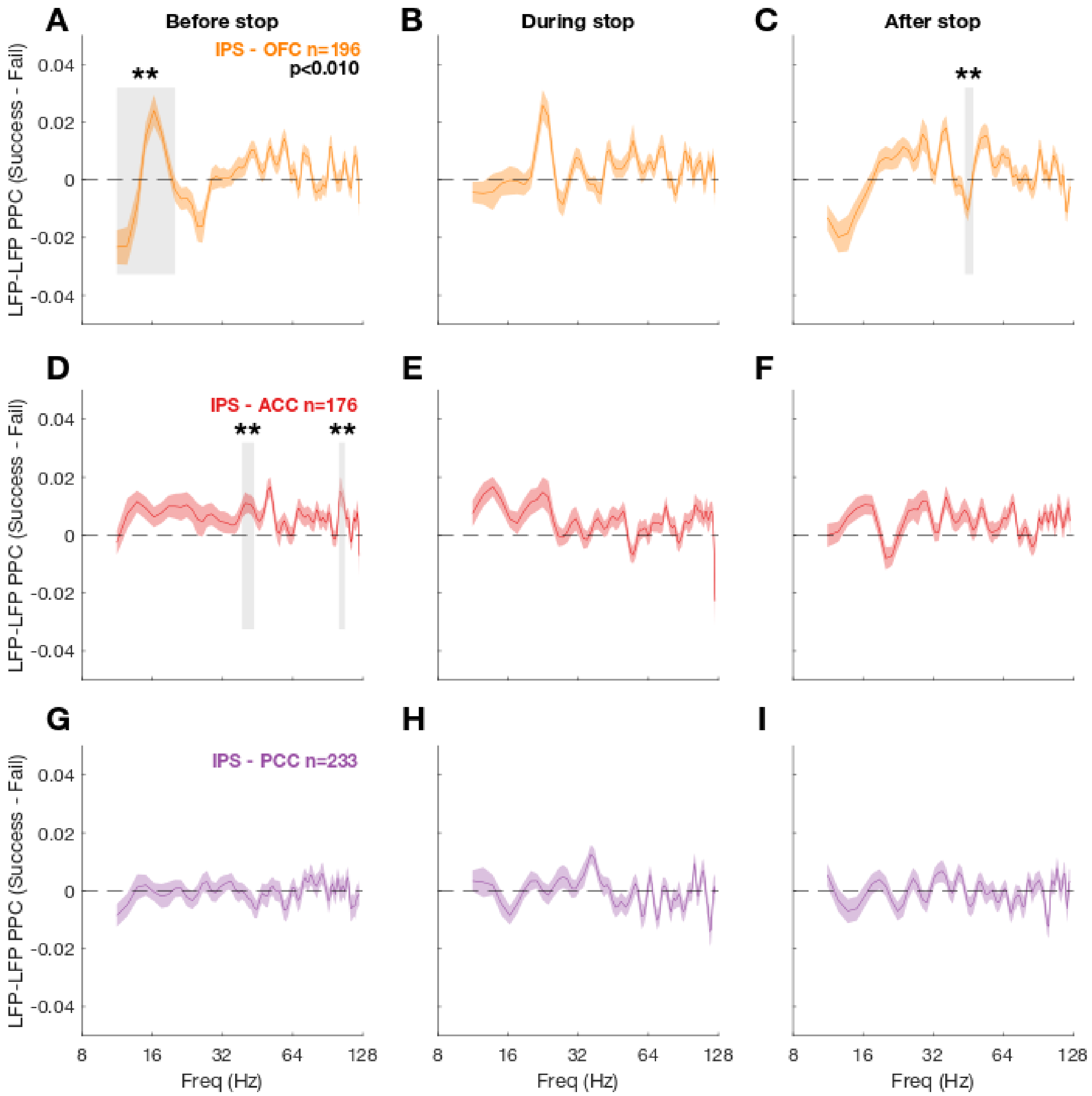


### Figure S7. IPS-OFC, IPS-ACC, IPS-PCC PPC analysis

Same format as Figure 5A-C. We measured LFP-LFP PPC between IPS and OFC (A-C), between IPS and ACC (D-F), and between IPS and PCC (G-I). Black asterisks and gray vertical shaded regions denote $p < 0.01$ (paired t-test at each frequency). No significant PPC modulations in (B) and (E-I). (A) There is significant modulation in PPC between IPS and OFC in 11-20 Hz (Before-stop) and (C) in 44-48 Hz (After-stop). (D) There is significant modulation in PPC between IPS and ACC in 39-44 Hz and 101-108 Hz (Before-stop).

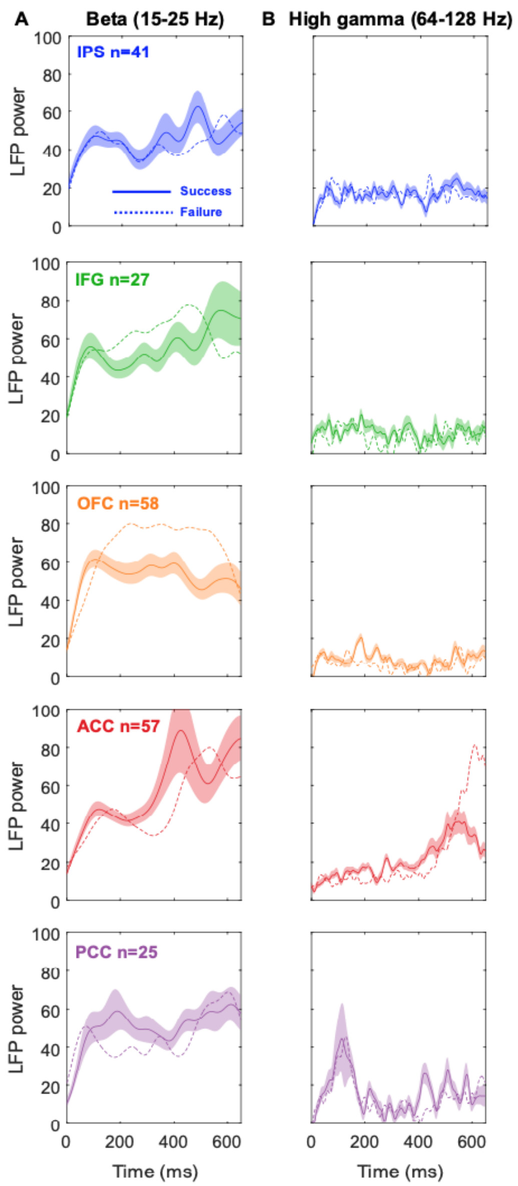


Figure S8. LFP power in beta (15-25 Hz) and gamma (64-128 Hz) frequency ranges aligned to the Stop-cue onset from 7 patients (#1, #4, #5, #7, #8, #9, #10)

(A) Beta LFP power in ACC IFG, IPS, OFC, PCC. Shaded regions denote ± SEM. Shaded regions for Stop-fail trials are omitted. (B) Gamma LFP power.
